# The Alzheimer’s disease protective P522R variant of *PLCG2*, consistently enhances stimulus-dependent PLCγ2 activation, depleting substrate and altering cell function

**DOI:** 10.1101/2020.04.27.059600

**Authors:** Emily Maguire, Georgina E. Menzies, Thomas Phillips, Michael Sasner, Harriet M. Williams, Magdalena A. Czubala, Neil Evans, Emma L Cope, Rebecca Sims, Gareth R. Howell, Emyr Lloyd-Evans, Julie Williams, Nicholas D. Allen, Philip R. Taylor

## Abstract

Recent genome-wide association studies of Alzheimer’s disease (AD) have identified variants implicating immune pathways in disease development. A rare coding variant of *PLCG2*, which encodes PLCγ2, shows a significant protective effect for AD (rs72824905, P522R, *P*=5.38×10^−10^, Odds Ratio = 0.68). Molecular dynamic modelling of the PLCγ2-R522 variant, situated within the auto-inhibitory domain of PLCγ2, suggests a structural change to the protein. Through CRISPR-engineering we have generated novel *PLCG2*-R522 harbouring human induced pluripotent cell lines (hiPSC) and a mouse knockin model, neither of which exhibits alterations in endogenous *PLCG2* expression. Mouse microglia and macrophages and hiPSC-derived microglia-like cells with the R522 mutation, all demonstrate a consistent non-redundant hyperfunctionality in the context of normal expression of other PLC isoforms. This signalling alteration manifests as enhanced cellular Ca^2+^ store release (∼20-40% increase) in response to physiologically-relevant stimuli (e.g. Fc receptor ligation and Aβ oligomers). This hyperfunctionality resulted in increased PIP_2_ depletion in the cells with the PLCγ2-R522 variant after exposure to stimuli and reduced basal detection of PIP_2_ levels *in vivo*. These PLCγ2-R522 associated abnormalities resulted in impairments to phagocytosis (fungal and bacterial particles) and enhanced endocytosis (Aβ oligomers and dextran). PLCγ2 sits downstream of disease relevant pathways, such as TREM2 and CSF1R and alterations in its activity, direct impacts cell function, which in the context of the inherent drugability of enzymes such as PLCγ2, raise the prospect of manipulation of PLCγ2 as a therapeutic target in Alzheimer’s Disease.

## Introduction

The gene encoding phospholipase C gamma 2 (PLCγ2) was recently linked by whole-exome microarray genome wide association (rs72824905/P522R, *P*=5.38×10^−10^) to Alzheimer’s disease (AD), the most common form of dementia (Qiu *et al*., 2009). The majority of AD cases are late-onset AD (LOAD) with a sporadic aetiology. The heritability of AD is estimated at 58-79% (Gatz *et al*., 2006), demonstrating a strong genetic influence on the development of AD. Genome wide association studies (GWAS) have identified genomic regions associated with LOAD (Harold *et al*., 2009, Hollingworth *et al*., 2011, Lambert *et* al., 2013, Sims *et al*., 2017, Sims *et al*., 2020). As well as these common variants, a number of rare genetic variants have been reported which implicate immunity and a crucial role for microglia in disease development. The rare protective R522 protein-coding variant in *PLCG2* is one such example (Sims *et al*., 2017). PLCγ2 is the first classically drug-targetable molecule identified from a LOAD genetic study. Understanding how this variant may lower the risk of developing AD is essential, not just for a greater understanding of disease mechanisms, but also for guiding future drug development approaches. Fundamental questions remain unresolved: How might the variant alter PLCγ2 function? Does the R522 variant manifest a functional alteration in physiologically-relevant cells (such as microglia)? When expressed at endogenous expression levels does this alteration occur at the level of cell signalling? Is its function non-redundant in the context of expression of other additional PLC isoforms as found in microglia? This study sought to address these questions.

PLCγ2 is a member of the Phospholipase C (PLC) family of enzymes. All 13 members of the PLC enzyme family play a role in signal transduction pathways (Kadamur and Ross, 2013) and there are two PLC gamma molecules, 1 and 2, which share a high sequence homology but differ greatly in their expression profiles. PLCγ2 is largely considered to be expressed in hematopoietic cells and is involved in the regulation of both the development and function of various hematopoietic cells (Koss *et al*., 2014). In the brain, PLCγ2 is expressed predominantly in microglia. PLCγ2 has been shown to catalyse the hydrolysis of phosphatidylinositol(4,5)bisphosphate [PI(4,5)P_2_] and this serves to increase concentrations of cytosolic facing diacylglycerol (DAG) and inositol 1,4,5 trisphosphate (IP_3_), which in turn increases the concentration of intracellular Ca^2+^ (Kadamur and Ross, 2013, Frandsen and Schousboe, 1993, Zheng *et al*., 2009). PLCγ enzymes have also been implicated in aberrant cellular responses linked to complex disease development (Bunney and Katan, 2010, Everett *et al*., 2009). One example is an association between dominantly inherited complex immune disorders and gain-of-function mutations in PLCγ2, such as deletions in the PLCG2-associated antibody deficiency and immune dysregulation (PLAID) disorder that occur in the cSH2 autoinhibitory domain (Ombrello *et al*., 2012).

The cSH2 domain of PLCγ2 prevents enzymatic activity and receptor ligation is believed to displace this domain revealing the active site, it is therefore not surprising that deletions here can confer increased enzymatic activity, in certain conditions (Milner, 2015). Through an ENU mutagenesis strategy, Yu *et al*. identified a gain of function variant of PLCγ2 which leads to subsequent hyperactivation of B cells and innate immune cells. They showed an autoimmune and inflammatory response, which implicates PLCγ2 as a key regulator in autoimmune and inflammatory disease (P. *et al*., 2005). In microglia, PLCγ2 regulates multiple signalling pathways leading to functions such as phagocytosis, secretion of cytokines and chemokines, cell survival and proliferation (Jay *et al*., 2015, Ulrich *et al*., 2014, Wang *et al*., 2015). With this in mind it is important to consider how PLCγ2 function may effect microglial responses in the context of amyloid deposition and other neuroinflammatory components of dementia seen in AD. Deficiency in TREM2, an upstream receptor that signals through PLCγ2, has been previously linked to increased risk of AD (Sims *et al*., 2020, Guerreiro *et al*., 2013). PLCγ2 also sits downstream of CSF1R, inhibitors or which are currently in trials in the context of Alzheimer’s disease. Thus, the role of PLCγ2 in Alzheimer’s disease may be complex.

The recently described rare coding R522 variant of PLCγ2 is a protective variant that causes a neutral to positive amino acid change. We hypothesised this mutation causes a structural and functional impact on the protein by altering the behaviour of the loop in which residue 522 is found and subsequently other protein domains. In this manuscript, through the use of specific CRISPR-mediated engineered alterations of human iPSC (hIPSC) and mice, we demonstrate that the R522 variant results in a hyperfunctional enzyme, which in AD relevant cells is associated with enhanced Ca^2+^-signalling, substrate depletion and altered phagocytic and endocytic responses. The relevance of this to protection from AD and the potential consequences for therapeutic approaches are discussed.

## Materials and Methods

### Mice

B6(SJL)-Plcg2em1Adiuj/J (#29598) mice were generated as detailed in the Supplementary Method section and are available from The Jackson Laboratory. The mice were maintained under specific pathogen free conditions with environmental enrichment as standard and all experiments were conducted in accordance with UK Home Office (Animal Scientific Procedures Act 1986) and Institutional guidelines.

### Mouse Microglia and Macrophage cells

Microglia and macrophages differentiated in M-CSF from conditionally-immortalized progenitors (M-MØP) were isolated and cultured as described in Supplementary methods from wildtype Plcg2^P522^ and variant Plcg2^R522^ knockin mice. Validation approaches are described in Supplementary methods

### Human Kolf2 hiPSC cell lines

*PLCG2* variant Kolf2 hiPSC cell lines were derived and differentiated into microglia as described in Supplementary methods. Validation of these cells are contained within the Supplementary methods

### GapmeR knock down of mouse Plcg2

Plcg2 knockdown was achieved in M-MØP cells using an antisense LNA GapmeR produced against mouse Plcg2 (Qiagen). Cells were grown to 80% confluence (∼2.4 × 105 cells per well) in 6 well plates. 3 µl of Liptoectamine RNAimax (Invitrogen) was diluted in 150 µl RPMI media and mixed with 150 pmol of siRNA in 150 µl RPMI media and left for 5 minutes. 250 µl of siRNA-lipid complex was added to cells in 3 ml RPMI and the cells were left for 48 hours. Cells were then washed and used for Ca^2+^ imaging (as described below) and knock down of Plcg2 was confirmed with qPCR (see Supplementary Methods). ΔΔCt values were used to calculate fold change and comparison of experimental and control samples used to measure knockdown. Negative and transfection controls were performed in parallel.

### Single cell imaging of Ca^2+^-signalling

Single cell Ca^2+^ imaging was performed using the ratiometric cytosolic Ca^2+^ probe Fura-2 (Barreto-Chang and Dolmetsch, 2009; Dolmetsch and Lewis, 1994(Lee *et al*., 2015)). Cells in chamber slides (Ibidi) were then loaded with 2 µM Fura2-AM (Abcam ab120873) solution at room temperature in the dark for 45 minutes. Cells were then washed and imaged in Ca^2+^-free media and EGTA was used to set the Fmin. Cells were imaged on an inverted Zeiss Colibri LED widefield fluorescence microscope with a high-speed monochrome charged coupled device Axiocam MRm camera and Axiovision 4.7 software. Primary mouse microglia and M-MØP cells were exposed at set time points to anti-FcγRII/III (2.4G2) antibody at 5 µg/ml (Stemcell technologies), LPS (50 ng/ml) (Sigma Aldrich) or oligomers of Aβ1-42 (40 µM)(preparation details in Supplementary Methods). This was followed by Ionomycin (2 µM) (Tocris) as a positive control and to set the Fmax. Cells were inhibited by pre exposure for 2 hours with Edelfosine (10 µM) (Tocris) or U73122 (5 µM) (Sigma). hIPSC-derived microglia were exposed to anti-CD32 (Fisher 16-0329-81, 5 µg/ml). The ratio between the excitation at 360 nm and 380 nm was used to indicate intracellular Ca^2+^ levels with regions of interest drawn over whole cells.

### Measurement of Ca^2+^-signalling by whole culture fluorimetry

Mouse macrophages and microglia were washed with HBSS then loaded with 2 µM Fluo8-AM solution (Stratech) at room temperature in the dark for 1 hour. Cells were then washed and imaged in Ca^2+^-free media and EGTA used as a negative control. Plates were warmed in a plate reader (Spectramax Gemini EM) to 37°C and Cells were exposed at set time points to anti-FcγRII/III, LPS or oligomers of Aβ1-42, as above. This was followed by Ionomycin (2 µM) as a positive control. Levels of fluorescence were detected at ex 490 nm/ em 520 nm (Hagen et al., 2012). Cells were inhibited by pre exposure for 2 hours with Edelfosine (10 µM) (Tocris) or U73122 (5 µM) (Sigma) or Xestospogin C (5 µM) (Abcam).

### Assays of phagocytosis and endocytosis

hIPSC-derived microglia, primary microglia, and M-MØP were prepared as described in supplementary methods. pHrodo-labelled-zymosan (Invitrogen, P35364) and -E. coli (Invitrogen, P35361) bioparticles were diluted to 0.25mg/ml and pHrodo-labelled dextran (10,000MW, Invitrogen, P10361) was diluted to 0.1mg/ml in live cell imaging solution (Invitrogen, P35364) then sonicated for 15 minutes using Biorupter sonicator (Diagenode). On the day of the assay, hiPSC-derived microglia were washed with 100µl of live cell imaging solution and a phase image (20X) was taken of the plate using an IncuCyte Zoom Live-Cell Analysis System (Essen). The live cell imaging solution was then removed, and a fresh 100µl was added prior to the addition of 25µl of the phagocytic/endocytic cargo. The hiPSC-derived microglia were put into the IncuCyte and imaged at 20X every 20 minutes for 4 hours. An analysis pipeline was then set up on the IncuCyte which provided both percentage confluence (from the first phase scan) and total red object integrated intensity for each well at each time point. Total red object integrated intensity per well per time point was then divided by percentage confluence in order to obtain a normalized fluorescence reading per well. Similarly, the primary microglia and M-MØP cells were then imaged using an EVOS FL Auto 2 onstage incubator system (Thermofisher). Images at x40 from cells in the 8 well chamber slide were taken every 30 minutes until the 2 hour mark, using the EVOS (transmitted light and RFP filter). Cells in the 96 well plate were read using the 544/590 filter on a BMG FLUOstar plate reader (BMG Labtech) to detect increased fluorescence per well. A BCA assay was run in parallel to normalize data to protein levels.

### Clearance of amyloid oligomers

hiPSC-derived microglia, primary microglia and M-MØP cells were prepared as described in Supplementary methods. Fluorescent amyloid oligomers (Eurogentech, AS-60479-01) were reconstituted to 0.1mM by adding 1% NH_4_OH followed by PBS. This stock was then diluted in astrocyte conditioned media (See Supplementary Methods) or RPMI (0.5µM) and was added to each well of the 96-well plate. After 2 hours, media was removed and replaced with 0.2mg/ml trypan blue for 2 minutes to quench extracellular fluorescence. The trypan blue was then removed and cells were washed twice in live cell imaging buffer. For iPSC-derived microglia green fluorescence was measured in an IncuCyte Zoom Live-Cell Analysis System. Total green object integrated intensity per well was divided by percentage confluence in order to obtain a normalized fluorescence reading per well.

Primary microglia and M-MØP cells green fluorescence was measured using the GFP filter on a BMG FLUOstar plate reader to detect increased fluorescence per well. Cells were imaged using the EVOS FL Auto 2 with the GFP filter. A BCA assay was run in parallel to normalize results to protein levels. Fluorescence was normalized further by taking the first reading at 0 mins as 0 arbitrary units (A.U.) for phagocytosis/endocytosis.

### Immunostaining of PIP2 *in vitro*

Microglia and M-MØP cells were prepared as described in Supplementary methods. Cells were then exposed to anti-FcγRII/III (2.4G2) antibody at 5 µg/ml, LPS (50 ng/ml) or oligomers of Aβ1-42 (40 µM). Designated wells were inhibited by pre exposure for 2 hours with Edelfosine (10 µM) or U73122 (5 µM). Other wells were pre exposed to LPS (50 ng/ml) (Sigma Aldrich) or oligomers of Aβ1-42 (40 µM) for 4 hours. At set time points cell were fixed (4% PFA,10 minutes) then washed with PBS twice. Cells were then permeabilized (0.05% saponin in PBS+tween-20 (0.01%), 10 minutes) prior to washing with PBS and blocking (60 minutes,10% BSA in PBS+tween-20 (0.01%)). Cells were then exposed to fluorescein conjugated Anti-PI(4,5)P2 IgM (Z-G045 Echelon Bioscience, 1:200, 4 hours). Cells were then washed three times with live cell imaging solution and green fluorescence was measured using the GFP filter on a BMG FLUOstar plate reader to detect increased fluorescence per well. Similarly cells were imaged using the EVOS FL Auto 2 with the GFP filter at x40 and intensity was measured with Image-Pro Premier software (MediaCybernetics). A BCA assay was run in parallel to normalize. Fluorescence was further normalized by taking the negative control unstained well at 0 mins as 0 A.U.

### Live cell assay of DAG *in vitro*

Microglia and M-MØP cells were prepared as described in Supplementary methods. DAG levels were detected using Green Down Assay from Molecular Montana (D0300G) following the manufacture instructions. Further details are provided in the Supplementary Methods.

### PIP isolation from cells *in vitro*

Microglia and M-MØP cells were prepared as described in Supplementary methods. Cells were then exposed to anti-FcγRII/III (2.4G2) antibody at 5 µg/ml or oligomers of Aβ1-42 (40 µM). Designated wells were inhibited by pre exposure for 2 hours with 3-a-aminocholestane (20µM, B-0341), LY294002 (10µM, B-0294) or SF1670 (5µM, B-0350) (Echelon Bioscience). Cells were then exposed to ice cold 0.5 M TCA (Trichloroacetic acid T6399 Sigma) for 5 minutes. Cells were then scraped off and spun (3000 RPM,7 minutes,4°C). The cell pellet was washed twice with 5% TCA/ 1 mM EDTA, vortexed for 30 seconds and centrifuged (3000 RPM, 5 minutes). The pellet is then vortexed twice with MeOH: CHCl3 (2:1) for 10 minutes at room temperature and centrifuged (3000 RPM,5 minutes). The pellet is then vortexed with MeOH: CHCl3:12 N HCl (80:40:1) for 25 minutes at room temperature and centrifuge at 3000 RPM for 5 minutes. The pellet is then discarded and CHCl3 and 0.1 N HCl are added. The tube is then centrifuged at 3000 RPM for 5 minutes to separate organic and aqueous phases. Collect the organic phase into a new vial and dry in a vacuum dryer for 60 minutes.

### Quantification of PIP isoforms by mass ELISA

PIP isoforms were quantified from extracted lipids using PI(4,5)P2 Mass ELISA (K-4500), PI(3,4)P2 Mass ELISA (K-3800) and PIP3 Mass ELISA (K-2500s) from Echelon Biosciences using manufacturer instructions.

### Immunohistochemical detection of PIP2 in mouse brain

Wildtype Plcg2^P522^ and variant Plcg2^R522^ knockin mice were aged to 2 months then trans-cardiac perfusion with 4% PFA was performed. The brains were removed and post fixed with 4% PFA for 4 hours at 4°C. Brains were then washed and cyroprotected with 30% sucrose until sunk. Brains were then embedded in OCT and cut using a CryoStar NX50 cryostat (Thermo Scientific) at 12µm. Brain sections were mounted on super adhesive slides (Lieca) and stored at -80°c prior to fixing (2% PFA,10 minutes). Slides were then permeabilized (0.1% saponin,PBS-T (0.1%),30 minutes) prior to quenching using 2mg/ml ammonium chloride and blocking (10% BSA and 5% goat serum in PBST). Cells were then stained overnight at 4°C with anti-PI(4,5)P2 IgM (Z-G045 Echelon Bioscience 1:100) and anti-IBA1 (013-26471 Alphalabs 1:200). Sections were stained with DAPI in mounting media (H-1200 Vectorlabs) and sealed with nail varnish. Images were taken at x63 using a Cell Observer spinning disc confocal (Zeiss). Position in the brain section were found using anatomical markers and images were taken at the primary somatosensory cortex and CA1 hippocampus. Images were analysed using Zeiss Blue 3.0 (Zeiss) and Image-Pro premier. Iba1 was used as a marker for microglia and used to create a region of interest. Inside that region of interest the level of green fluorescence caused by staining of PIP2. Intensity of fluorescence per cell area was calculated and background fluorescence was subtracted.

### Molecular dynamic modelling of PLCγ2 variants

No complete protein databank (PDB) structure was available for the PLCγ2 protein. To overcome this, homology modelling was used to construct the protein in its entirety using the I-TASSER server (Yang et al., 2015). The I-TASSER server uses a protein threading method to create a homology model and a model with a high C-score, a measure of protein quality, was selected and further scrutinised for quality using PROCHECK (Laskowski and MacArthur, 1993). The mutated version of the protein was created using the modify protein function in Discovery studio. Further details are provided in the Supplementary Methods.

### Quantification and statistical analysis

Statistical analysis were conducted using Prism (GraphPad) or R and its inbuilt ‘stats’ package. The details of the tests used and data representation are provided in the main text, in the results and figure legends sections.

## Results

### Molecular Dynamic predictions and structural impact of P522R variant

The HoPE server was used to report on the possible effects that the change in amino acid may have on the protein structure, and make some functional predictions (Venselaar *et al*., 2010). There are a number of obvious changes; to begin the mutated amino acid is bigger than the WT (Fig. 1A), it also carries a positive charge and is less hydrophobic than the WT. The size change is predicted to cause an interaction between this amino acid and other parts of the protein, this interaction will cause a knock-on effect on protein structure. HoPE also shows the sPH domain, where the mutation is found, to interact with two other domains which are involved in protein function.

**Figure 1.**
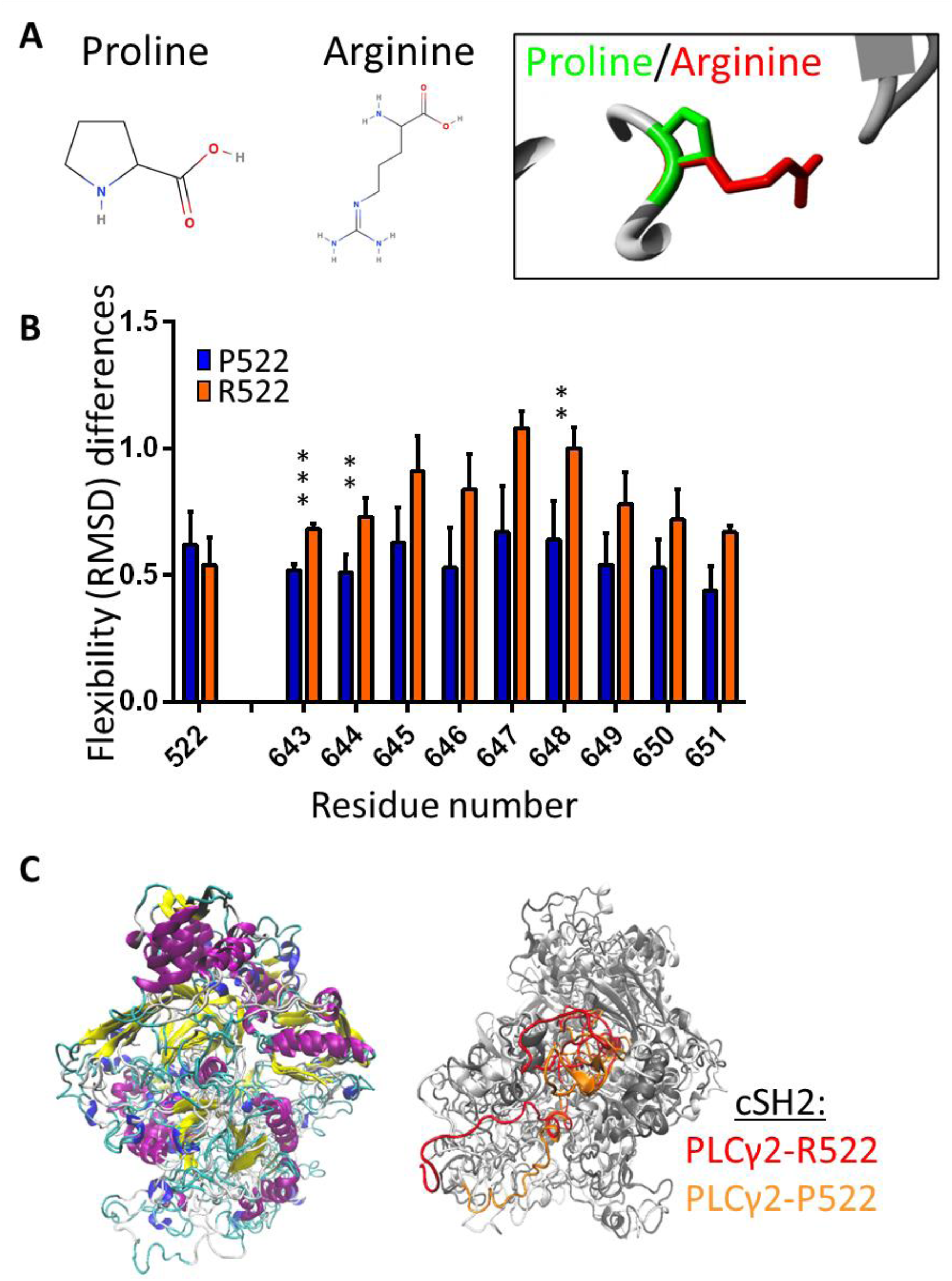
Molecular Dynamic and structure of P522R variant. **A**. 1. Proline structure, 2. Arginine structure, 3. The mutation site image, produced by the HoPE server, depicting the P522 in green with the R522 mutation in red (Venselaar et al., 2010). **B**. Flexibility (RMSD) differences between the wild type and mutated protein, significant differences are highlighted in the figure. RMSD was measured per residue over the whole simulation time using the GROMACS tool g_rms and here averages were plotted with standard deviations represented in the error bars. Data were analysed using a Mann-Whitney U test in the R statistical package. **C**. 1. Wild type and Mutated structures overlaid using the RMSD alignment function in VMD, both are coloured using secondary structure colouring and overlap of domains can be seen. 2. This is the same overlaid structured viewed from an alternative side of the protein. In this image most of the structure is light grey with the cSH2 domain highlighted in orange (P522) and red (R522).

Both the WT and mutated protein simulations have consistent energy, pressure and volume outputs with very low standard errors, showing a good stability in the simulation. The root mean standard deviation (RMSD) can be used to record the overall flexibility, or that of individual amino acids. RMSD for the WT and p.P522R simulations are not statistically different (p=1.83e-09) indicating the mutation does not have an overall effect on the flexibility of the protein. However, mutation appears to have a greater effect on the flexibility of amino acid 522, along with other small areas of the protein including a section of a nearby loop containing residues 643-651 (Fig. 1B), which makes up part of the cSH2 domain. A number of these residues show statistically significant differences in flexibility (i.e. p<=0.05) between the WT and the mutated protein.

The p.522R mutation has a wider impact on the PLCγ2 protein than just the changing of the local loop structure in which it is found. The cSH2 domain is also significantly altered, in both position and structure, where the rest of the structure is conserved (Fig. 1). When compared to the wild type the mutated cSH2 domain is shifted to the right and now contains an alpha helix where before there was a random coil (Fig. 1C). There is also a loss of two alpha helices from the WT to the mutated protein. Further to this change, hydrogen bonding is increased when proline is switched to an arginine, with an increased number of donors and acceptors. Proline has one acceptor and arginine has two acceptors and five hydrogen bonding donor sites. During the simulation, WT p.P522 formed just two hydrogen bonds with two different near-by residues, 512, and 527. This suggests that the WT p.522 residue has local interactions only. The mutated p.R522 residue, however, formed 17 hydrogen bonds with six different residues, including the glycine amino acid at residue 948 (Supplementary Table 1). The 948 residue is found within the PLC-Y-Box – part of the TIM barrel binding domain, and this change in interaction causes a previously randomly coiled section of the protein to form a helix (Figure 1). The change in flexibility and position of the cSH2 domain could feasibly modulate PLCγ2’s ability to trigger intracellular Ca^2+^release.

### The R522 variant of Plcg2 confers increased receptor-mediated Ca^2+^ release in macrophages and microglia

Four independent conditionally-immortalised macrophage precursor (MØP) polyclonal cell lines were derived from both male and female *Plcg2*^P522^ and *Plcg2*^R522^ mice. The genotype of donor mice was confirmed prior to use (Supplementary Fig. 1A,B) and similar expression of *Plcg2* in M-CSF-differentiated ‘M-MØP’ macrophages of the two genotypes was verified by qPCR (Supplementary Fig. 1C).

Results from Fura2 intracellular Ca^2+^ assays demonstrated that M-MØPs differentiated from *Plcg2*^R522^ precursor cell lines produced a significantly greater rise in intracellular Ca^2+^ after exposure to anti-FcγRII/III (Fig. 2A), compared to their Plcg2^P522^ expressing counterparts. As such the R522 variant appears to be hyper functional compared to WT in this *in vitro* system. All the cell lines produced a similar response after exposure to Ionomycin. This suggests both cell lines were alive and capable of Ca^2+^ response.

**Figure 2.**
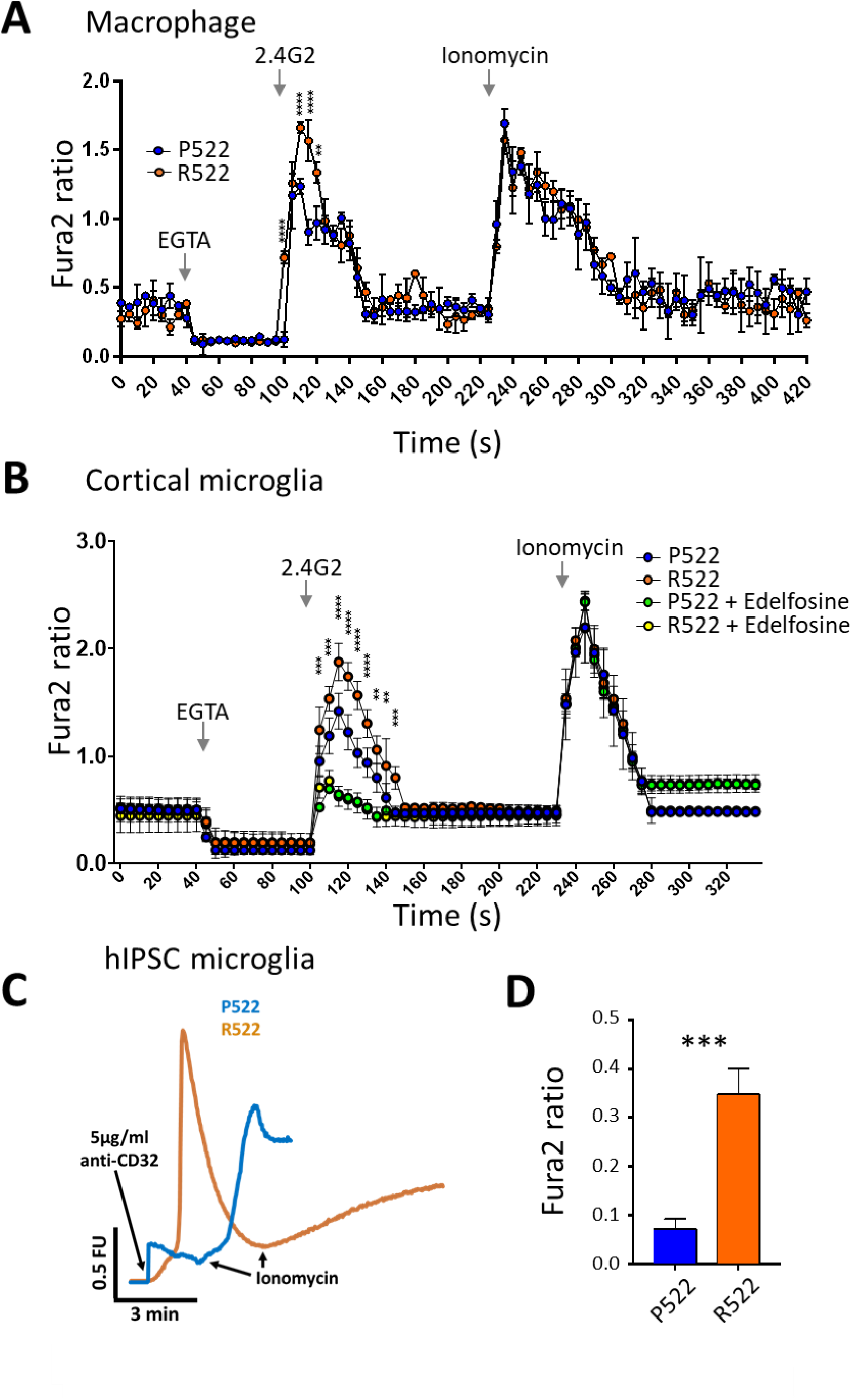
Receptor-mediated release of Ca^2+^ in macrophages and microglia. **A**. Fura2 340/380 time traces from M-CSF-differentiated macrophages derived from conditionally-immortalised macrophage precursor cell lines (M-MØP) of *Plcg2*^P522^ mice (P522, blue) and *Plcg2*^R522^ mice (R522, red). One set of cell lines, generated from male mice, is shown. Cells were exposed to 5µg/ml anti-FcγRII/III along with 20µM EGTA and 2µM Ionomycin as indicated. **B**. Fura2 340/380 time traces from primary microglia derived from the cortex of *Plcg2*^R522^ mice (blue: P522) and *Plcg2*^R522^ mice (red: R522) with or without pre-exposure for 2 hours with Edelfosine (10µM). Cells were exposed to 5µg/ml anti-FcγRII/III along with EGTA and 2µM Ionomycin. Data in A and B shows the mean±SD of 3 independent experiments analysed by two-way ANOVA with Sidak post-hoc tests. *PLCG2*^R522^ hiPSC-derived microglia show increased cytosolic Ca^2+^ influx in comparison to controls following activation of PLCγ2 using anti-CD32. Cytoplasmic Ca2+ increase following activation of PLCγ2 using anti-CD32 Representative Ca^2+^ traces (**C**) and graphical summary (**D**). 3 independent *PLCG2*^R522^CRISPR-engineered clones were examined. Data shown is mean±SD of 4 independent experiments and were analysed by one-way ANOVA with Tukey’s multiple comparison test (** = p<0.01; *** = p<0.001; **** = p< 0.0001).

To confirm the R522 variant is also hyper functional in primary cells, microglia were cultured from the cortex and hippocampus of neonatal *Plcg2*^P522^ and *Plcg2*^R522^ mice. Using a Fura2 assay it was shown that anti-FcγRII/III exposure resulted in a greater Ca^2+^ response in the R522 variant expressing cells when compared to the WT. This was found in microglia cultured from both the cortex (Fig. 2B) and hippocampus (Supplementary Fig. 1D). The Ca^2+^ response in both regional types was inhibited using Edelfosine suggesting the Ca^2+^ response was due to the function of PLCG.

We next repeated Fura2 intracellular Ca^2+^ assays in microglia-like cells derived from either control PLCγ2^R522^ or variant PLCγ2^R522^ expressing hiPSCs to confirm our findings with human cells (Fig 2. C,D). Similar to the mouse cells, activation of PLCγ2 using anti-CD32 (anti-FcγRII) resulted in a greater cytosolic Ca^2+^ increase in PLCγ2^R522^ variant cells compared to control PLCγ2^P522^ cells.

In order to confirm that the assays were measuring PLCγ2-induced Ca^2+^ release a number of approaches were taken, employing both Fura2 and Fluo-8 assays. First, PLCγ was inhibited with Edelfosine and U71322 and the IP3 receptor blocked with Xestospongin c during stimulation of macrophages with anti-FcγRII/III (2.4G2) (Fig. 3A). All 3 inhibitors prevented the FcγR-induced Ca^2+^ response (Fig. 3A). To confirm that the intracellular Ca^2+^ release was due to PLCγ2 activity in this assay, GapmeRs were used to knockdown *Plcg2* mRNA in mouse macrophages, prior to study (Fig3. B). GapmeR-mediated knockdown was very effective (Fig3B, left panel) and prevented FcγRII/III-mediated Ca^2+^ release (Fig. 3B, right panel). Similar results were obtained with primary microglia (Fig. 3C). Next, we examined the Ca^2+^ response of primary microglia to stimulation with LPS (50ng/mL) and Aβ_1-42_ oligomers (40µM) (Fig. 3D). In both cases, cells expressing the R522 variant exhibited enhanced Ca^2+^ response compared to WT.

**Figure 3.**
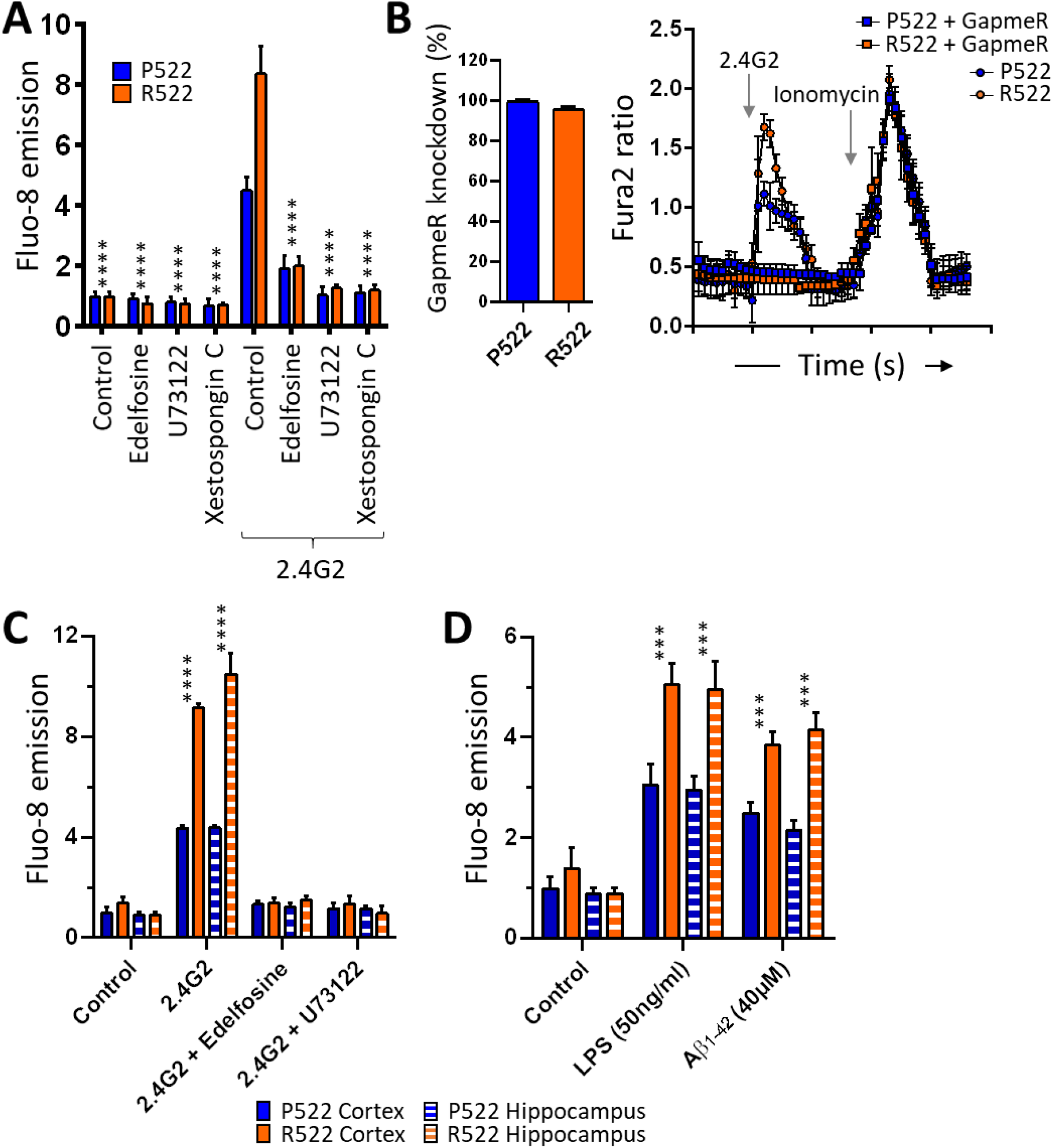
Specificity of PLCγ2 in response of macrophages/microglia to physiologically-relevant stimuli. **A**. M-MØP (blue: P522; red: R522) were loaded with Fluo-8 Ca^2+^ indicator and examined for peak changes in fluoresce after exposure to 5ug/ml anti-FcγRII/III (2.4G2) with or without pre-exposure for 2 hours with Edelfosine (10µM) or U73122 (5µM) or Xestospogin C (5µM). **B**. Left panel shows LNA GapmeR percentage knockdown of *Plcg2* gene expression in M-MOP cells within the P522 control (blue) and R522 (red) variant lines. Right panel shows Fura2 340/380 time traces from M-MØP cell lines (blue: P522; red: R522) which have undergone *Plcg2* knockdown (square symbols) using an antisense LNA GapmeR. Cells were exposed to 5µg/ml anti-FcγRII/III and 2µM Ionomycin. **C**. Microglia from *Plcg2*^P522^ (blue: P522) mice and *Plcg2*^R522^ mice (red: R522) from neonate cortex (solid colour) or hippocampus (striped) were loaded with Fluo-8 Ca^2+^ indicator. These cells were then examined for peak changes in fluoresce after exposure to 5µg/ml anti-FcγRII/III. These readings were taken with or without pre-exposure for 2 hours with Edelfosine (10µM) or U73122 (5µM). **D** Cortical microglia from *Plcg2*^P522^ (blue: P522) mice and *Plcg2*^R522^ mice (red: R522) from neonate cortex (solid colour) or hippocampus (striped) were loaded with Fluo-8 Ca^2+^ indicator. These cells were then examined for peak changes in fluoresce after exposure to LPS (50ng/ml) or Aβ_1-42_ oligomers (40µM). Data shown as the mean±SD of 3 independent experiments. Data in A, C and D were analysed by two-way ANOVA with Dunnett’s multiple Comparison test. See also Supplementary figure 3.

### Expression of the AD protective PLCγ2 R522 variant impaired phagocytic and enhanced endocytic clearance

Phagocytosis can occur via a variety of mechanisms and provides a key function of microglia and macrophages within the brain (Janda *et al*., 2018). Furthermore, PLCγ2 is downstream of TREM2 and CSF1R, both known to regulate aspects of the cell biology of these lineages, including microglial phagocytosis (Xing *et al*., 2015). We assessed how the R522 variant of PLCγ2 influenced phagocytic activity using the standardised pHrodo-labelled bioparticles derived from *E*.*coli* and Zymosan. Mouse macrophages, microglia and human iPSC derived microglial (Fig. 4A-C, respectively) all exhibited significantly reduced phagocytosis of *E*.*coli* when expressing the R522 variant compared to the common P522 variant. Similar results were obtained with all three cell types with zymosan particles (Fig4. D-F).

**Figure 4.**
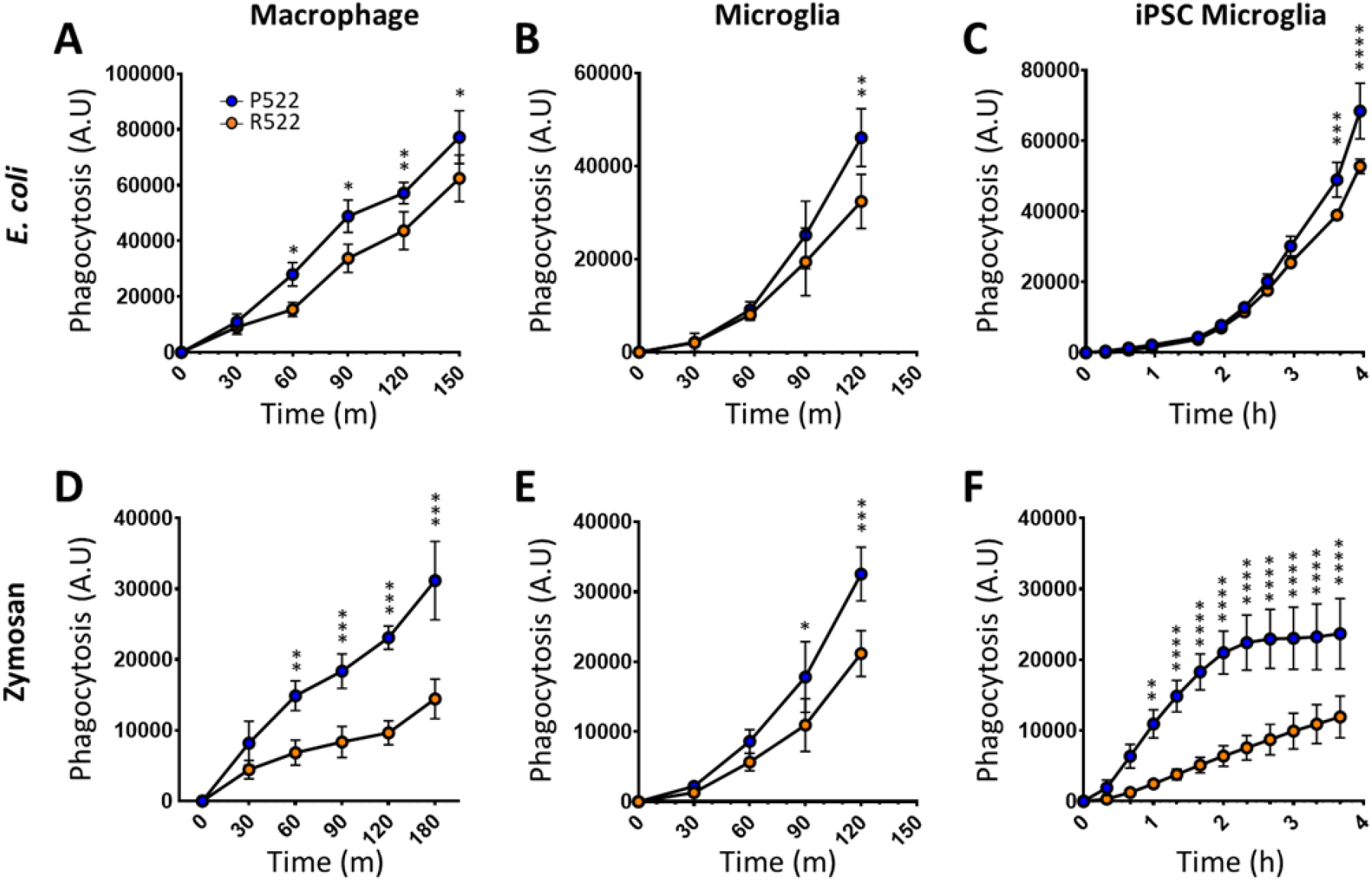
Protective R522 M-MOP and microglia display decreased phagocytosis of *E. coli* and zymosan when compared with common P522 variant cells. Phagocytotic activity of R522 and P522 M-MOP and microglia was assessed using pHrodo Red *E. coli* and zymosan BioParticles. Phagocytosis arbitrary units (A.U) describes the amount of bioparticle cellular fluorescence emission at each time point. *E*.*coli* uptake in M-MOP **(A)**, primary mouse microglia **(B)**, hiPSC-derived microglia **(C)**. Zymosan uptake in M-MOP (D), primary mouse microglia **(E)** and hiPSC-derived microglia **(F)**. .For hIPSC-derived microglia, 3 isogenic P522 and 3 isogenic R522 clones were examined with at least 6 wells in 3 independent experiments. All microglia and M-MOP data shows the mean±SD of 3 independent experiments were analysed by two-way ANOVA using Sidak multiple comparison test. *=p<0.05, **=p<0.01, ***=p<0.001, ****=p<0.0001. (blue: P522; red: R522)

Subsequently, we examined the endocytic clearance of fluorescent Aβ_1-42_ oligomers by the same cells (Fig5. A-C). In all cases, expression of the R522 variant resulted in enhanced clearance of the oligomers. Similar results were obtained with a model cargo (pHrodo-labelled 10KDa Dextran) (Fig.5. D-E).

**Figure 5.**
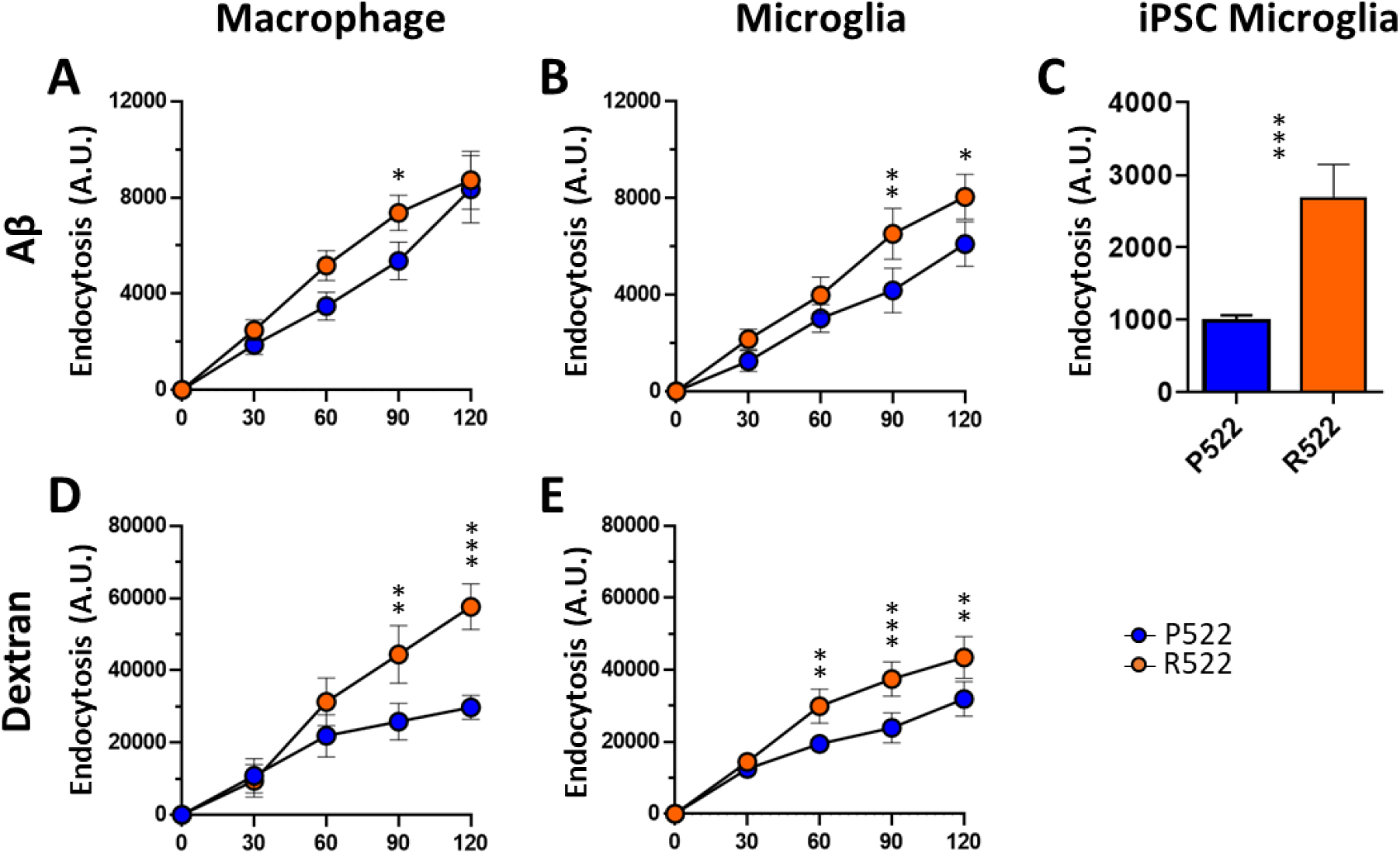
Protective R522 M-MOP and microglia display increased endocytosis of soluble Aβ_1-42_ oligomers and dextran when compared with common P522 variant cells. Endocytic activity of R522 and P522 M-MOP and microglia was assessed using FITC soluble Aβ_1-42_ oligomers and pHrodo Red Dexran (10,000 MW). Endocytosis arbitrary units (A.U) describes the amount of bioparticle cellular fluorescence emission at each time point. Aβ_1-42_ oligomer uptake in M-MOP **(A)**, primary microglia **(B)**, and hiPSC-derived microglia **(C)**. Dextran uptake in M-MOP **(D)** and primary microglia **(E)**. For hiPSC-derived microglia, 3 isogenic P522 and 3 isogenic R522 clones were examined with at least 6 wells in 3 independent experiments. All microglia and M-MOP data shows the mean±SD of 3 independent experiments were analysed by two-way ANOVA using Sidak’s multiple comparison test. *=p<0.05, **=p<0.01, ***=p<0.001. (blue: P522; red: R522)

### *Plcg2*^R522^ expressing cells exhibit greater reduction of PIP2 levels after PLCγ_2_ activation

We next examined the change in levels of the PLCγ2 substrate PIP_2_ during exposure to anti-FcγRII/III, LPS (50ng/mL) or Aβ_1-42_ oligomers (40µM) by immunofluorescence (Fig6A, Supplementary Figure 2). These stimuli caused a significant reduction in PIP_2_ levels in both macrophages and microglia. Cells expressing *Plcg2*^R522^ demonstrated a greater reduction of PIP_2_ compared to the *Plcg2*^P522^ controls. This reduction in PIP_2_ was prevented using the inhibitors edelfosine or U71322. This was also demonstrated in pre incubation with LPS (50ng/mL) and Aβ_1-42_ oligomers (40µM) but only over longer time frames (Fig 6A,B). Using a DAG fluorescent sensor we similarly saw that DAG levels increase after exposure to anti-FcγRII/III, LPS (50ng/mL) and Aβ_1-42_ oligomers (40µM) (Fig. 6C,D). Similarly both the *Plcg2*^R522^ expressing cells demonstrated a greater increase in DAG compared to the *Plcg2*^P522^ controls (Fig. 6C,D). This increase was prevented using the inhibitors edelfosine or U71322. Using a PI(4,5)P_2_ mass ELISA to detect a more specific change in PIP species we confirmed the reduction in PI(4,5)P_2_ after exposure to anti-FcγRII/III and that there was a significantly greater decrease in the *Plcg2*^R522^ expressing cells compared to the control (Fig. 7A). Using 3-a-aminocholestane (SHIP1 Inhibitor), LY294002 (PI3-K Inhibitor) or SF1670 (PTEN Inhibitor) did not prevent this reduction. Mass ELISA for PI(3,4)P_2_ demonstrated no significant difference between the two variants after exposure to anti-FcγRII/III with or without the inhibitors (Fig. 7B). Quantification of PI(3,4,5)P_3_ by mass ELISA demonstrated no significant difference between the two variants after exposure to anti-FcγRII/III. But addition of either a SHIP1 inhibitor or a PTEN inhibitor with anti-FcγRII/III resulted in a significant reduction in PI(3,4,5)P_3_ consumption (Fig. 7C).

**Figure 6.**
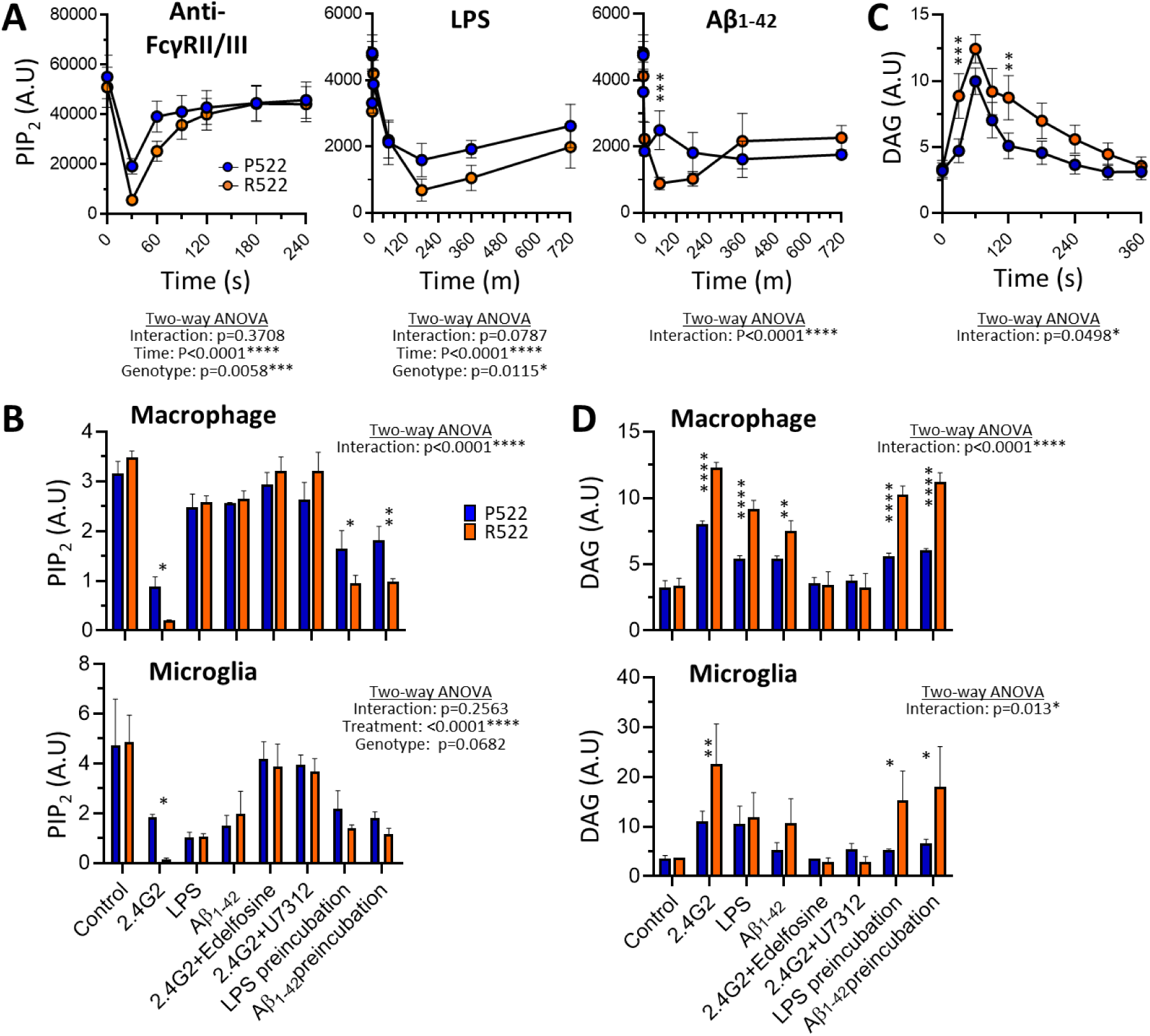
Protective R522 M-MOP and microglia display increased depletion of PIP_2_ after exposure to physiologically-relevant stimuli compared to P522 M-MOP and microglia. The level of PIP_2_ in M-MOPS was examined by measuring immunofluorescence from images at set time points after exposure after exposure to 5µg/ml anti-FcγRII/III (2.4G2), 50 ng/ml LPS or oligomers of 40 µM Aβ1-42 **(A)**. The levels of PIP_2_ was also measured using a plate reader after exposure to physiologically relevant stimuli in M-MOPS and primary mouse microglia **(B)**. DAG level were measured using a live cell assay in M-MOPS (and primary mouse microglia as a time course from immunofluorescence **(C)** and plate reader after exposure to physiologically relevant stimuli **(D)**. Data shows the mean±SD of 3 independent experiments were analysed by 2 way ANOVA with Sidak’s multiple comparison *=p<0.05, **=p<0.01, ***=p<0.001, ****=p<0.0001. (blue: P522; red: R522). See also supplementary figure 2.

**Figure 7:**
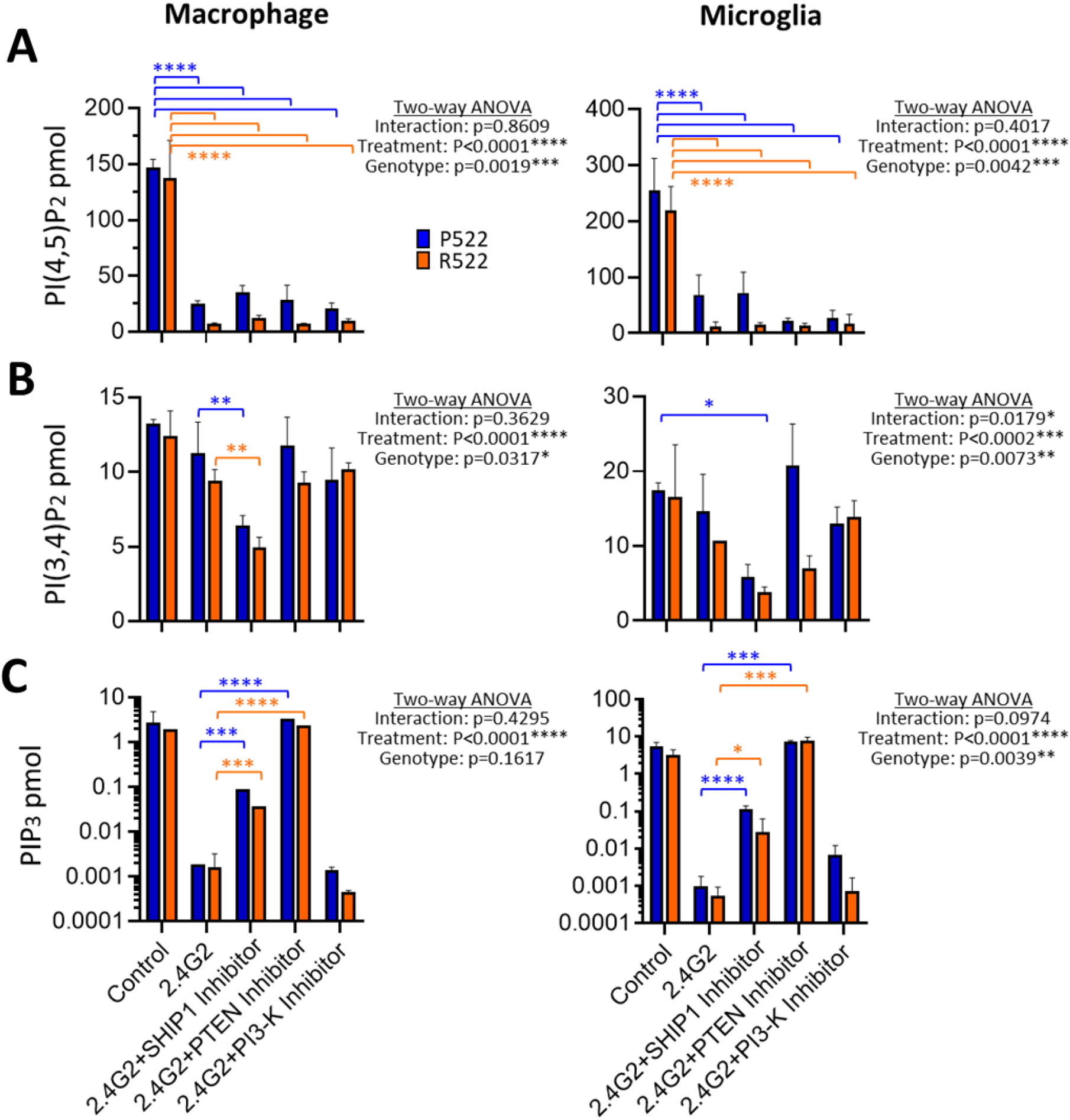
Protective R522 M-MOP and microglia display increased depletion of PI(4,5)P_2_ without corresponding compensation from other PIP species. A mass ELISA was used to detect specific PIP species after exposure to anti-FcγRII/III (2.4G2) with or without 3-a-aminocholestane (SHIP 1 inhibitor), SF1670 (PTEN inhibitor) and LY294002 (PI-3K inhibitor). In M-MOP cells (left panel) and primary microglia (right panel) PI(4,5)P_2_ **(A)**, PI(3,4)P_2_ **(B)** and PIP_3_ **(C)** were detected. All data shows the mean±SD of 3 independent experiments were analysed by 2 way ANOVA (log-transformed data was used in C) with Sidak’s multiple comparison tests performed *=p<0.05, **=p<0.01, ***=p<0.001, ****=p<0.0001. (blue: P522; red: R522)

### In vivo basal levels of PIP2 are reduced in cortical and hippocampal microglia in *Plcg2*^R522^ mice

The cortex and hippocampus of two month old *Plcg2*^R522^ and *Plcg2*^P522^ mice were examined for PIP_2_ levels in microglia by immunofluorescent staining for PIP_2_ and Iba1 (Fig. 8). Results were calculated from three separate groups of mice per variant. Representative immunostaining is shown in Fig 8A. In both the cortex and hippocampus (Fig. 8B) the levels of PIP_2_ in microglia were reduced by approximately 60% in the *Plcg2*^R522^ mice compared to the *Plcg2*^P522^ mice indicating a reduced basal level of PIP_2_ in AD vulnerable regions

**Figure 8.**
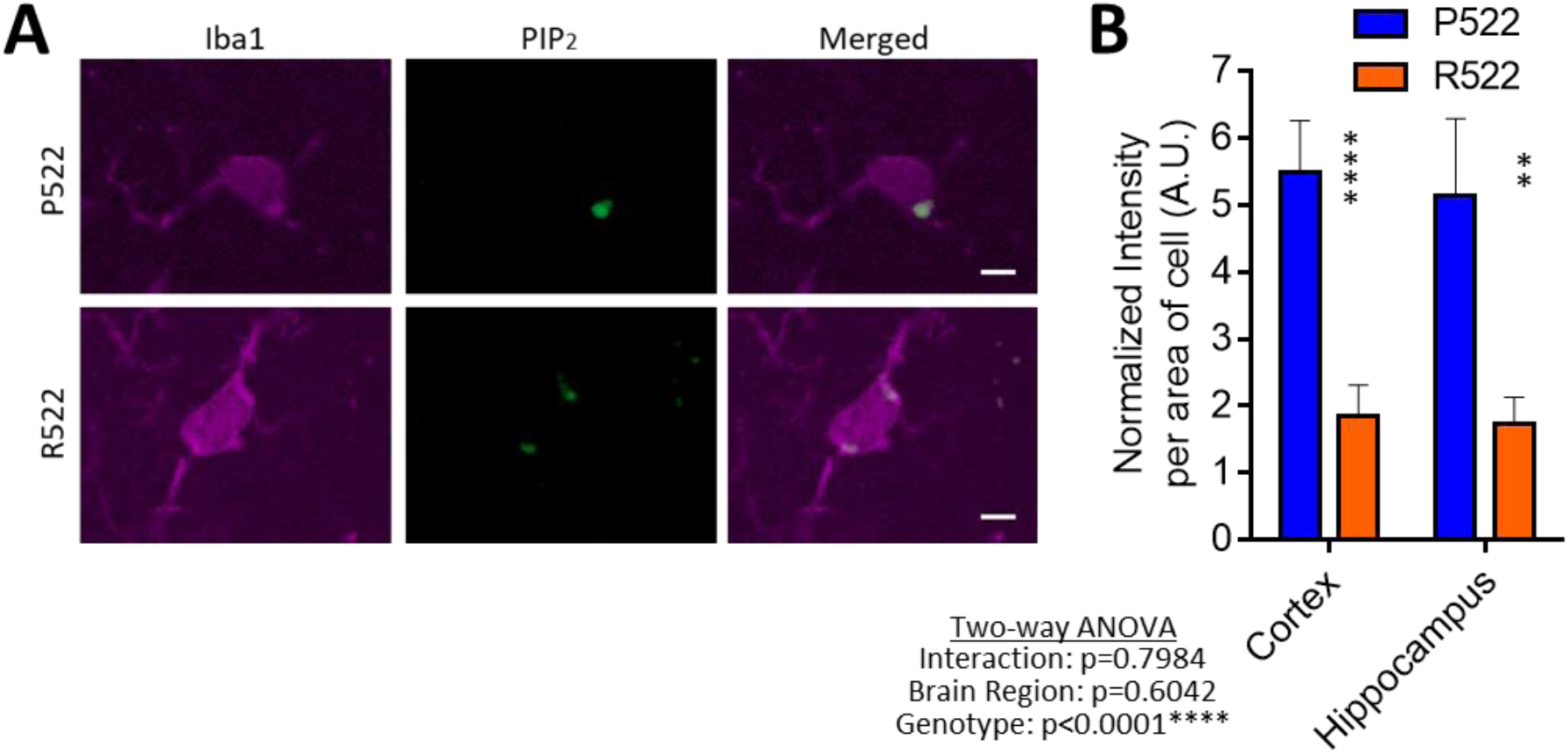
In vivo basal PIP_2_ levels are decreased in microglia of *Plcg2*^R522^ mice compared to *Plcg2*^P522^ mice. Cyrosections of *Plcg2*^R522^ and *Plcg2*^P522^ mouse brains were immunostained for Iba1 and PIP2 and images were taken in the cortex and hippocampus. Representative images of the cortex are displayed from *Plcg2*^P522^ and *Plcg2*^R522^ Iba1=Red and PIP_2_=Green, scale bar=5µm (**A**). Iba1 was used as a marker for microglia and PIP_2_ levels were detected by measuring the intensity of florescence per cell. The average fluorescence was calculated above background in the cortex and hippocampus (**B**). All data shows the mean±SD of 3 independent experiments analysed using 2 way ANOVA with Sidak’s multiple comparison tests performed **=p<0.01, ****=p<0.0001. See also supplementary figure 4.

## Discussion

The heritability of AD is estimated to be between 58-79% (Gatz *et al*., 2006). This strong genetic influence in AD has provided novel insights into the mechanisms underlying disease and will hopefully lead to potential new approaches for drug design. One such opportunity is presented by the recent discovery of a protective rare coding mutation in *PLCG2* P522R; (Sims *et al*., 2017) PLCγ2 is an enzyme amenable to drug development approaches and hence presents an opportunity for intervention in the development of AD. To aid rational drug design and understand how PLCγ2 contributes to the disease process, it is important to understand how rare coding variants affect cell activation in myeloid cells and microglia, the primary cell expressing PLCγ2 in the brain. To address the role of the protective R522 mutation in PLCγ2 and enable modelling in a physiologically-relevant context, we have developed two novel models for its study: Kolf2 CRISPR gene-edited hiPSC and CRISPR gene-edited mice, both of which harbour the R522 variant, which is conserved across the species. The generation of these original models for the study of PLCγ2 and AD is an essential first step in understanding the underlying mechanisms that lead to disease.

We present evidence that the mutated R522 PLCγ2 protective variant is hyper functional and results in consistently increased intracellular Ca^2+^-signalling in response to a variety of receptor mediated stimuli in microglia and macrophages. This altered signalling likely directly contributes to protection against AD as it acts downstream of receptors, such as TREM2 and CSF1R, implicated in AD. An earlier study had indicated that the R522 variant may be hyperfunctional in COS7 and HEK293T expression systems (Magno *et al*., 2019). Importantly, our studies show that PLCγ2 with the R522 variant exhibits increased function in both human and mouse disease-relevant microglia when expressed at physiological normal levels, in the context of other PLC enzymes.

Computational modelling suggests that this hyperfunction may occur via inhibition of an auto-inhibitory domain within PLCγ2. Analysis revealed a change in flexibility and position of the cSH2 domain which could feasibly modulate PLCγ2’s ability to indirectly trigger intracellular Ca^2+^release. Increased hydrogen bonding may have important implications for protein function as it shows a change in orientation and location of the mutated residue loop. This could have a considerable structural impact on a binding domain and it is possible that a change in the binding domain leads to the observed movement of the cSH2 domain to which it is connected. The cSH2 domain of this protein is believed to play a critical role in stabilising the early signalling complex which is stimulated by BCR crosslinking, where the domain inhibits the active site of PLCγ2 (Wang *et al*., 2014). A known gain of function mutation (G993) in PLCγ2 increases external Ca^2+^ entry causing an auto-immune and inflammatory response (P. *et al*., 2005). The cSH2 domain interacts with residues surrounding the catalytic active site, resulting in the auto-inhibition of the PLCγ enzymes. This inhibition is reversed by the interaction of cSH2 with a phosphorylated tyrosine residue, this domain movement allows for access to the active site. PI(3,4,5)P3, binds to the active site and recruits PLCγ isoforms to the plasma membrane, resulting in their activation. Studies in PLCß2 have shown residues in the region surrounding the loop, which connects two parts of the catalytic domain to limit substrate access to the active site. If the same is true of PLCγ2 then the R522 mutation may be causing a similar change to auto-inhibition (Everett *et al*., 2009). These functional predictions help explain our observations of increased PLCγ2 activity within R522 variant microglia and macrophages.

Having observed increased PLCγ2-mediated IP3-depenent Ca^2+^ signalling within our models and devised a possible mechanism underlying this hyper-function at the protein level, we then investigated the effect of this hyper-functionality on PLCγ2’s substrate (PI(4,5)P_2_) and product (DAG). PI(4,5)P_2_ levels in R522 and P522 M-MOP were examined using fluorescein conjugated Anti-PI(4,5)P_2_ following addition of either anti-FcγRII/III antibody (to stimulate PLCγ2), LPS, or Aβ. Following addition of anti-FcγRII/III antibody, we found that the protective R522 variant resulted in a significantly greater drop in PI(4,5)P_2_ when compared to P522 control at 30 seconds, and this drop then appears to take longer to return to baseline. LPS and Aβ addition also appears to induce a greater drop in PI(4,5)P_2_ in the R522 compared to P522 expressing cells (Fig 6A). Observations of reduced PI(4,5)P_2_ are further supported in Fig 6B where anti-PI(4,5)P_2_ staining (measured via plate reader assay) in M-MOP and microglia, following activation with anti-FcγRII/III antibody resulted in decreased PI(4,5)P_2_ in the protective R522 compared with P522.

PLCγ2 is positioned downstream of important signalling receptors, such as TREM2 and CSF1R and as such, would be expected to be undergoing tonic activation. With the observation that stimulation of PLCγ2 leads to enhanced depletion of its substrate (PI(4,5)P_2_) by the protective R522 variant, compared to the common P522 allele, we hypothesized that the R522 variant may lead to a state of depressed substrate levels *in vivo*. We addressed this by assessing PI(4,5)P_2_ levels in microglia *in situ* in the brains of young P522 and R522 expressing mice. We observed a marked reduction in microglial PI(4,5)P_2_ in the R522 mice compared with their controls (Fig 8) demonstrating that the impact of the protective variant was evident in healthy young animals.

Live cell DAG assays demonstrated that protective R522 M-MOP and microglia produce more DAG than P522 controls when stimulated with anti-FcγRII/III antibody (Fig 6F-H). Moreover DAG levels remained higher overtime in the R522 cells, which is consistent with a faster diffusion profile of DAG compared with PI(4,5)P_2_) (Xu *et al*., 2017). Both direct addition and pre-treatment of M-MOP with LPS or Aβ again resulted in more DAG production in the R522 expressing cells compared with P522 cells, and a similar pattern of response was observed in the microglia.

PI(4,5)P_2_, PI(3,4)P_2_ and PIP3 were also measured via mass ELISA in P522 and R522 M-MOP and primary microglia, confirming reduced PI(4,5)P_2_ in the protective R522 variant following PLCγ2 activation with anti-FcγRII/III antibody. This effect was not prevented by co-treatment with SHIP1, PI3K, or PTEN inhibitors, demonstrating PI(4,5)P_2_ reduction as an upstream event of these enzymes following PLCγ2 activation.

Taken together, our observations demonstrate increased PLCγ2 activity in R522 variant expressing cells when compared to those expressing P522 resulting in increased PI(4,5)P_2_ consumption, and consequent DAG production. The increased PI(4,5)P_2_ utilisation associated with the R522 variant coupled with a slow recovery of PI(4,5)P_2_ levels would lead to a depleted PI(4,5)P_2_ pool compared with common P522 variant cells. The delayed return to baseline, tonic activation of PLCγ2 and consequent reduction in available PI(4,5)P_2_ levels could limit further activation of PLCγ2 within this time, potentially preventing chronic enzyme activation or reducing sensitivity to further receptor-mediated stimulation. Aligned to these observations, mice expressing PLCγ2^R522^ exhibited reduced microglial PI(4,5)P_2_ *in vivo*.

To understand how hyperactivity of PLCγ2 affects cell behaviour, we first investigated phagocytosis by the P522 and R522 expressing microglia and macrophages. By measuring uptake of pHrodo-labelled *E*.*coli* and zymosan particles. We observed an overall reduction in the levels of phagocytosis in all R522 expressing models when compared to the common P522 variant expressing controls. This phenotype could be attributed to reduced PI(4,5)P_2_ levels in the R522 variant cells. PI(4,5)P_2_ levels are critical to modulation of actin dynamics as reduction of this lipid at the plasma membrane has been demonstrated to result in reduce phagocytosis in RAW 264.7 cells (Botelho *et al*., 2000). This appears to be due to PI(4,5)P_2_ roles in actin filament formation: a key step in phagocytosis (Scott *et al*., 2005). Several recent findings suggest that increased phagocytosis may be an aggravating factor in AD (Nizami *et al*., 2019). For one, microglial activation and subsequent phagocytosis of stressed but still viable neurons, has been postulated to enhance AD pathology (Gabandé-Rodríguez *et al*., 2020). Increased phagocytosis of apoptotic neurons has been observed in APOE4 overexpressing cells (Muth *et* al., 2019), and inhibiting microglial phagocytosis appears to prevent inflammatory neuronal death (Neher *et al*., 2011). Furthermore, increased synaptic pruning, which occurs via microglial phagocytosis of synapses, has also been implicated in AD (Rajendran and Paolicelli, 2018). Reduced phagocytic activity in the R522 variant microglia and subsequent reduction in phagocytosis of damaged but viable neurons and synapses may contribute to the protective effect assigned to the R522 variant of PLCγ2.

We also assessed endocytic clearance of 10kDa dextran and soluble Aβ_1-42_ oligomers, both of which are endocytosed via micropinocytosis (Wesén *et al*., 2017). We observed increased uptake in all R522 models when compared to the common P522 variant. Reasons for this increased uptake by cells expressing the protective variant are currently unclear, although several studies have previously implicated PI(4,5)P2 in this process (Brown *et al*., 2001). Soluble Aβ has been demonstrated to perturb metabolic processes, induce the release of deleterious reactive compounds, reduce blood flow, inhibit angiogenesis, and induce mitochondrial apoptotic activity (Watson *et al*., 2005). Enhanced internalization of Aβ_1-42_ with R522 variant microglia when compared with common P522 could therefore reduce some of these toxic effects. Together, the results from our phagocytosis and endocytosis assays, which can be in part explained by observations of reduced PI(4,5)P_2_, suggest mechanisms by which the R522 variant may protect against AD.

In summary we have shown a consistent increase in enzymatic activation in novel human and mouse models due to the AD protective R522 mutation in PLCγ2. This is associated with depletion of PI(4,5)P_2_ both *in vitro* and *in vivo* and reduced phagocytic and increased endocytic clearance. These alterations in cell activities could directly impact on clearance of damaged cells and synapses *in vivo* both during disease and under homeostatic conditions where tonic activation of PLCγ2 would be expected. Reduced PI(4,5)P_2_ in the protective R522 variant expressing microglia, whilst allowing for short bursts of PLCγ2 activity, may limit such longer term enzyme activation. The generation of these original models for the study of PLCγ2 and AD has provided novel insight into the regulation of cell function and is an essential first step in understanding the underlying mechanisms that lead to disease.

## Acknowledgments

This work was supported by the UK Dementia Research Institute at Cardiff, Dementia Platform UK and Centre for Ageing and Dementia Research. The Moondance Foundation, P.R.T. is also supported by a Wellcome Trust Investigator Award (107964/Z/15/Z). G.E.M. is supported by a Ser Cymru II Fellowship, which is part funded by the European Regional Development Fund though the Welsh Government. J.W. is supported by Innovative Medicines Initiative (115736) and the Medical Research Council UK (HQR00720, MR/K013041/1). The work was in part supported by Eisai Inc. Additional costs were supported through an ARUK pump priming award (ARUK-NC2017-WAL), an ARUK Collaboration grant (ARUK-IRG2015-7 to E.L.E.) and MRC Partnership Award (MR/N013255/1 to N.D.A.). MODEL-AD is funded by U54 AG054345. The mouse CRISPR project was performed by The Jackson Laboratory Genetic Engineering Technologies group. This work was also funded in part by AG055104 (M.S. and G.R.H.).

## Author contributions

G.E.M., T.P. and E.M contributed to acquisition and interpretation of data and to the drafting of the work. H.M.W., M.A.C., N.E., E.L.C., contribute to the acquisition and analysis of data. M.S. and G.R.H contributed to the design of the work and interpretation of the data. R.S., E.L-E., J.W., N.D.A and P.R.T contributed to conception and design of the work and E.L-E, J.W., N.D.A and P.R.T contributed to the drafting of the work. All authors approved the submission.

