## Supplementary Materials for "The Alzheimer’s disease protective P522R variant of *PLCG2*, consistently enhances stimulus-dependent PLCγ2 activation, depleting substrate and altering cell function"

### Supplementary figures and tables

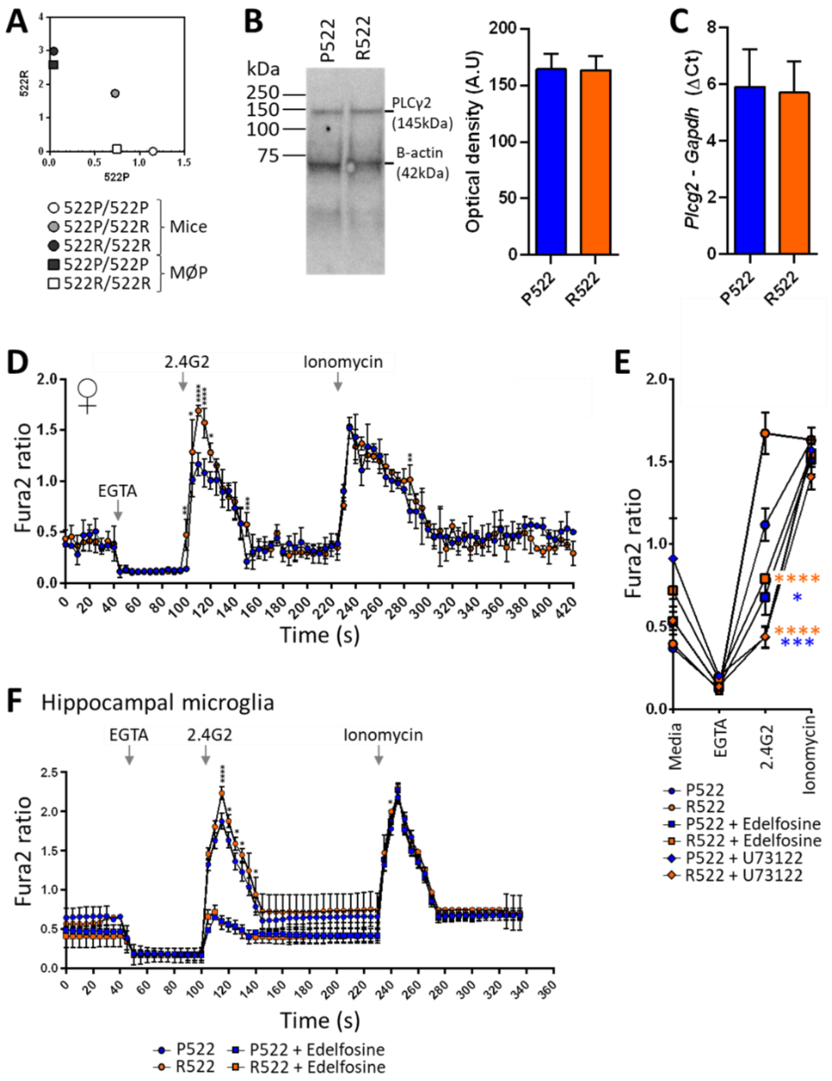


**Figure S1. Related to Figure 2,3. Novel R522 knockin mice and validation of mouse cells.**
qPCR endpoint analysis of genotype of *Plcg2*$^{P522}$ and *Plcg2*$^{R522}$ mice and conditional-immortalised macrophage precursor cells generated from them. **B.** Western blotting (representative image shown) of PLCγ2 in macrophage precursor cells with the P522 (blue) control and R522 (red) variants. Graph contains data from three repeats optical density normalised to loading control. **C.** qPCR results of gene expression of *Plcg2* in M-MØP cells

with the P522 (blue) control and R522 (red) variants. Results normalised to delta change of GADPH housekeeping gene expression. **D.** Fura2 340/380 time traces from M-CSF-differentiated macrophages derived from conditionally-immortalised macrophage precursor cell lines (M-MØP) of macrophages derived from the bone marrow cells of B6 wild type (P522, blue) mice and *Plcg2*$^{R522}$ mice (R522, red). Replicate independent cell lines generated from female mice are shown. Cells were exposed to 5ug/ml anti-FcγRII/III along with EGTA and 2uM Ionomycin as indicated. The data were analysed by two-way ANOVA with Sidak post-tests. **E.** Fura2 340/380 peaks of time traces from M-MØP (blue: P522; red: R522) treated with PLCγ-inhibitors. Cells were exposed to 5µg/ml anti-FcγRII/III along with EGTA and 2µM Ionomycin with or without pre-exposure for 2 hours with Edelfosine (10µM) or U73122 (5µM). Data were analysed by two-way ANOVA with Tukey's multiple comparison test. All graphical data represents mean±SD of 3 independent experiments, except D, where n = 4. **F.** Fura2 340/380 time traces from primary microglia derived from the hippocampus of neonate mice. Cells from B6 wild type (blue: P522) mice and *Plcg2*$^{R522}$ mice (red: R522) with or without pre-exposure for 2 hours with Edelfosine (10µM). Cells were exposed to 5ug/ml anti-FcγRII/III along with EGTA and 2uM Ionomycin. The data were analysed by two-way ANOVA with Sidak's post-tests. Data represents mean±SD of 3 independent experiments.

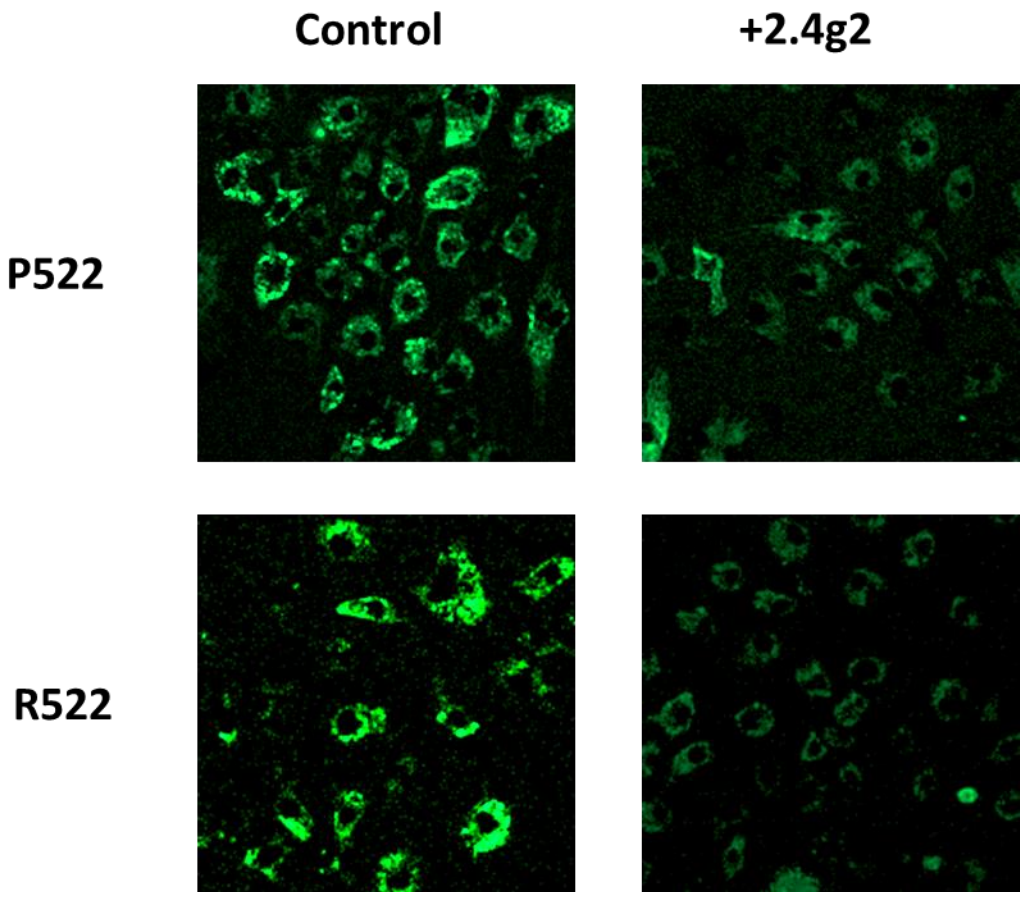


**Supplementary figure 2: Representative images of PIP2 staining in mouse macrophages.** Images taken of mouse macrophages from the *Plcg2*$^{P522}$ and *Plcg2*$^{R522}$ cell line stained against PIP2 (green) with (right panel) or without (left panel) 2.4G2 exposure

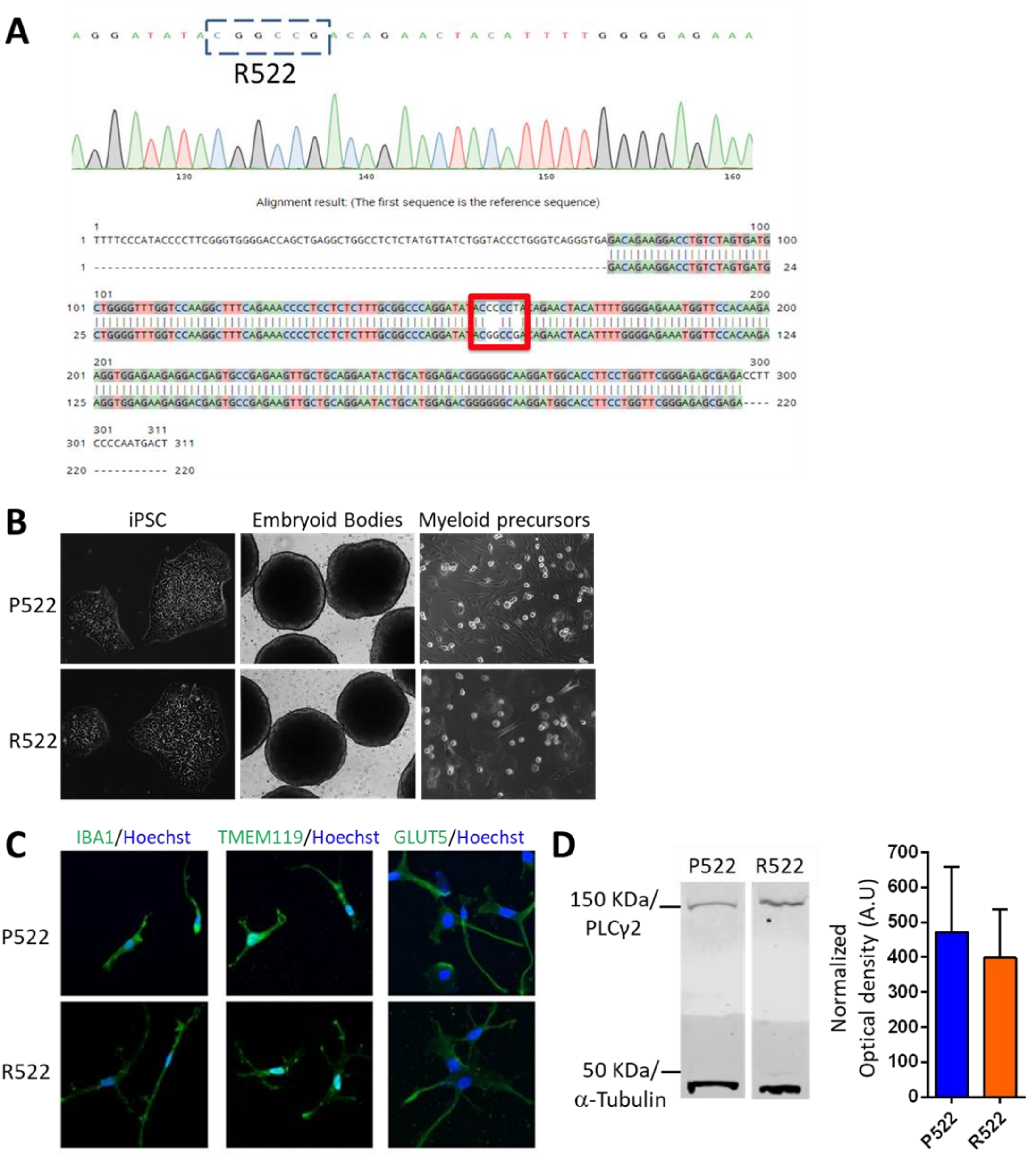


**Supplementary figure 3 Related to Figure 3. CRISPR-engineered R522 allele in human iPSC cells.**

Isogenic *PLCG2*$^{R522}$ variant iPSC lines were generated using CRISPR-assisted gene editing. Guide-RNAs were combined with Cas9 (HiFi Cas9 nuclease V3) to induce a double stranded break (DSB) in the specific P522 site in *PLCG2*. A P522R oligo was also introduced as a template for DNA repair enzymes. This resulted in the introduction of the mutation and the generation of 9 homozygotes and 7 heterozygotes clones for the R522 variant mutation within the KOLF2 iPSCs. Sequencing confirmed introduction of the P522R mutation (Example shown). **B.** Differentiation of P522R variant iPSCs first to embryoid bodies (EB's) then to myeloid precursors closely mimics the differentiation of control iPSCs. **C.** Immunofluorescent staining indicates a microglial-like phenotype with cells of both lines expressing microglial markers (IBA1, TMEM119 and GLUT5). **D.** Western blotting (representative images shown, left) of PLCγ2 in myeloid precursor cells with the P522 (blue) control and R522 (red) variants quantified (right). Graph contains data from three repeats optical density normalised to loading control, data represents mean±SD.

**A** **B**

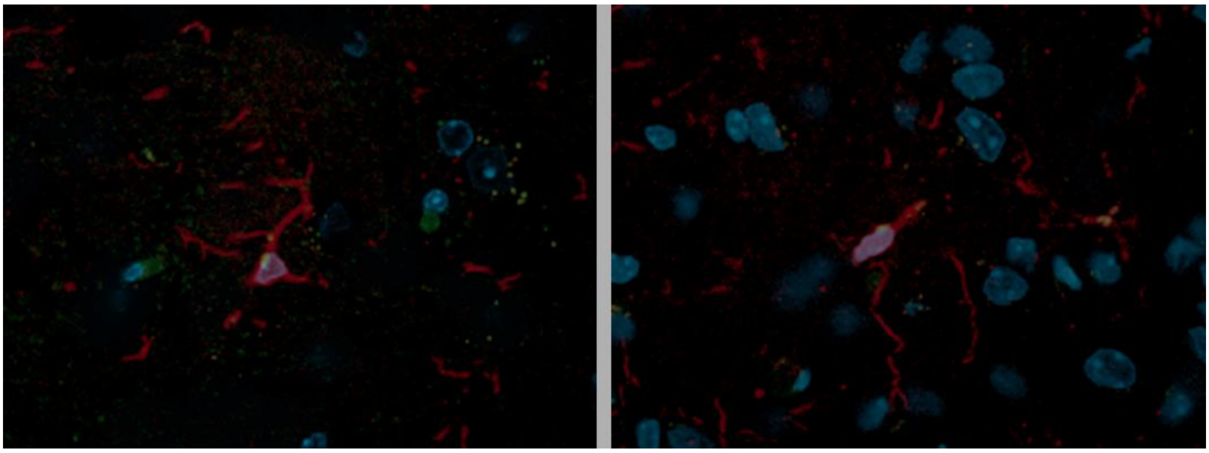

**Supplementary figure 4: Representative images of microglia in the cortex.** Images taken in the cortex of *Plcg2*$^{P522}$ (A) and *Plcg2*$^{R522}$ (B) mice labelled with Iba1 (red), PIP2 (green) and DAPI (blue).

**Supplementary tables:**

Table 1 - Hydrogen Bonding

| **WT/P522R** | **Donor** | **Acceptor** |
|---|---|---|
| WT | T512 | P522 |
| WT | H527 | P522 |
| P522R | T512 | R522 |
| P522R | Q519 | R522<br>(two sites) |
| P522R | R522 | E150 |
| P522R | R522 | P518 |
| P522R | R522 | L526 |
| P522R | R522 | H527<br>(three sites) |
| P522R | R522 | Q519<br>(two sites) |
| P522R | R522 | L526 |
| P522R | R522 | E948<br>(four sites) |
| P522R | R522 | E515 |

Table 2 – Wild-type and mutated protein simulation stability outputs

| Energy | Average | Standard Err. | Units. |
|---|---|---|---|
| | Wild-type Simulation | | |
| Total Energy | -3.62145e+06 | 190 | (kJ/mol) |
| Pressure | 1.01891 | 0.059 | (bar) |
| Volume | 3671.58 | 0.09 | (nm^3) |
| | Mutated Simulation | | |
| Total Energy | -3.62112e+06 | 190 | (kJ/mol) |
| Pressure | 0.960886 | 0.031 | (bar) |
| Volume | 3671.18 | 0.066 | (nm^3) |

Table 2 - Root mean square deviation (RMSD)

| | Wild Type | p.P522R Mutation |
|---|---|---|
| Mean | 0.49 | 0.50 |
| Standard Deviation | 0.07 | 0.07 |

### Supplementary methods

#### Generation of Plcg2 knock-in mice

Plcg2R522 knock-in mice (henceforth referred to as Plcg2R522) were generated by CRISPR/Cas9-assisted gene targeting in B6(SJL)-Apoetm1.1(Apoe*4)Adiuj/J (JAX#27894) mice as previously described. Briefly, microinjection mix contained 100ng of Cas9 and 50 ng of CRISPR guide (CCAAAATGCAGCTCCGTGGG) and 20ng of the 81nucleotide donor: (GAGCTGTAATAAGCCCTTTCGGATGCTTGTGGCTCAGGACACTCGCCCCACGGAGCTGCATTTTGGGGAGAAATGGTTCCACA) was used to engineer a C to G mutation (underlined) at nucleotide 1565 and change the CCC (proline) codon to CGC (arginine). This mutation disrupted the PAM sequence to prevent re-cutting. Founders were genotyped by PCR using forward primer 5'-GCGCTGCATGTTCCCTCTA-3' with reverse primer 5'-GGGCGTTACCAGAAGGAGAG-3' to generate a 375bp product, which was sequenced to identify the mutation. Positive founders were bred to in C57BL/6J (JAX #664) for two generations to breed away from the APOE4 allele. Once the line was established, genotyping was performed via Endpoint analysis using the amplifying primers: 5'-AATAAGCCCTTTCGGATGCT-3' and 3'-TCCGCACTGGTCCTACTCTC-5' with Wild Type Hex labelled probe 5'-Hex-AGGACACTCCCCCCACG - Black Hole Quencher 1-3' and Mutant 6-Fam labelled probe 5'-6-Fam-AGGACACTCGCCCCACG – Black Hole Quencher 1 – 3'. This model is available from the Jackson Laboratory as B6(SJL)-Plcg2em1Adiuj/J (#29598). Control (Plcg2P522) mice for the Plcg2R522 mice were generated from the littermates of the founders.

#### Mouse Plcg2-P522R variant genotyping

Ear biopsies were digested in 50 µl of mammalian lysis buffer (100 mM Tris-HCl pH 8.5, 5 mM EDTA, 0.2 % (w/v) SDS, 200 mM NaCl) containing Proteinase K at 100 µg/ml and incubated at a temperature of 56 °C whilst shaking at 1200 rpm for one hour in a 1.5 ml micro-centrifuge tube. Once dissolved, the sample was heated for a further 30 minutes at 72 °C to denature the Proteinase K. Once cooled to room temperature the sample was diluted by the addition of 450 µl of nuclease free water. The sample was frozen at -20 °C or immediately used for the qPCR reaction. Genotyping was performed in a 10 µl reaction volume incorporating 5 µl of the TaqMan® Fast Advanced Master Mix (Applied Biosystems), with 0.9 µM of each amplifying primer (see above), 0.25 µM of each probe (see above) and 4.5 µl of the sample digest both from above.

#### Generation and culture of conditionally-immortalised macrophage precursor (MØP) cell lines

Conditionally-immortalised macrophage precursor (MØP) were generated from bone marrow of wildtype Plcg2P522 and variant Plcg2R522 knockin mice and differentiated into macrophages with M-CSF (M-MØP) as previously described (Rosas et al., 2008; Rosas et al., 2011). Both male and female lines were established and the sex of the lines used in specific experiments are indicated in the main text. Briefly, CD117-enriched bone marrow cells (using CD117-biotin, BD Biosciences, and anti-biotin-MACS, Miltenyi Biotec) were infected with an estrogen-dependent Hoxb8 encoding pMX-IPs retroviral vector for conditional-immortalization and cultured in RPMI 1640 medium (Thermo Fisher Scientific) containing 10% (v/v) heat-inactivated fetal calf serum (FCS) (PAA Laboratories), 1%

penicillin/streptomycin (Pen/Strep), β-estradiol (1 μM) (Sigma-Aldrich), GM-CSF (10 ng/ml) (ImmunoTools), and puromycin (20 μM/ml) (Roth), After 10 days, puromycin was removed from the culture medium and the cells were passaged in the presence of β-estradiol and GM-CSF. When required, Plcg2P522 and Plcg2P522 MØP cells were further differentiated to M-MØP by application of 20 ng/ml M-CSF for 4 days in the absence of β-estradiol and GM-CSF.  For all experiments, M-MØP cells were grown to 80% confluence then starved of M-CSF for 4 hours prior to performing assays.

**Generation of primary mouse microglial cultures**

Primary neonate mouse microglia cultures were isolated from mixed glial cultures via a modified version of the previously described shaking technique (Tamashiro et al., 2012). Briefly, brains from Plcg2P522 and Plcg2R522 mice at P7-8 were collected and dissected into cortical and hippocampal sections in dissection media (HBSS with 0.1M HEPES (Thermo Fisher Scientific), 1x Penicillin/Streptomycin, 2mM L-Glutamine, 33 mM glucose (Sigma-Aldrich)).  Cells were enzymatically (0.05% Trypsin) and mechanically dissociated and seeded in on Poly-L Lysine (1 mg/ml Sigma Aldrich) coated T75 flasks in DMEM (Thermo Fisher Scientific) containing 10% (v/v) heat-inactivated FCS, 1x Penicillin/Streptomycin, 2mM L-Glutamine and M-CSF (10 ng/ml).  After 10-14 days cells were confluent and microglia were isolated from primary mixed cells via shaking at 200 rpm for 1.5 hour at 37 ºC. Microglia were plated in DMEM containing 10% heat-inactivated FCS, 1x Penicillin/Streptomycin, 2mM L-Glutamine and M-CSF (10 ng/ml) on 8 well chamber slides or 96 well plates and left overnight to attach. Purity of cell cultures was verified by immunostaining with anti-glial fibrillary acidic protein (Abcam ab7260) and anti-Iba1 (Merck MABN92).   For all experiments, primary microglia were grown to 80% confluence then starved of M-CSF for 4 hours prior to performing assays.

**Kolf2 hiPSC cell lines**

Isogenic PLCG2 P522/P522 and PLCG2 R522/R522 hiPSC clones were derived from the male parent Kolf2 cell line, previously validated for CRISPR genome editing; for availability and full genetic profile of Kolf2 cells see (http://www.hipsci.org/lines/#/lines/HPSI0114i-kolf_2). Kolf2 were grown on 10 μg/ml Geltrex coated culture plates (Nunc) in E8-Flex medium (ThermoFisher).  Colonies were passaged 1:10 after dissociating to cell clumps using Cell Dissociation Buffer (ThermoFisher).

**hiPSC culture and genome editing**

Kolf2 hiPSC clones harbouring the PLCG2R522 variant were generated by CRISPR gene editing. Deskgen CRISPR design tools (www.deskgen.com) were used to select a guide RNA (CCAAAATGTAGTTCTGTAGG) and ss-oligonucleotide homology directed repair template (5'GTCAGGGTGAGACAGAAGGACCTGTCTAGTGATGCTGGGGTTTGGTCCAAGGCTTTCAGAAACCCCTCCTCTCTTTGCGGCCCAGGATATACGGCCGACAGAACTACATTTTGGGGAGAAATGGTTCCACAAG3’). 2 nmol guideRNA and 20 nmol ATTO™ 550 labelled Alt-R® CRISPR-Cas9 tracrRNA were complexed in IDT buffer (all CRISPR reagents were purchased from Integrated DNA Technologies). The RNP complex was formed immediately prior to nucleofection by mixing the crRNA:tracrRNA complex with Alt-R® S.p. HiFi Cas9 Nuclease V3 (see Supplemtary Fig 3).

hiPSC cultures pretreated for 1 hour with 10 µM Y-27632 were dissociated into single-cell suspension using Accutase (Sigma). 1 × 106 cells were resuspended in Amaxa P3 nucleofection buffer (Lonza), mixed with Cas9 RNP complex and 100 pmol of ssDNA oligonucleotide repair template and nucleofected using program CA137 on the Amaxa-4D nucleofector. After replating in E8-Flex medium containing 10µM Y-27632 and overnight culture, nucleofected cells were harvested as a single cell suspension using Accutase and the brightest Atto550 labelled cells (~5000 cell) were harvested and replated onto a 10µg/ml Geltrex coated 10 cm plate. Cells were fed with E8-Flex medium containing 10 µM Y-27632 and 100 U/mL penicillin/streptomycin for the first 3days followed by E8-Flex only thereafter. Colonies derived from single cells were manually picked on day 7 into 96 well plates. 96 well plates were replica plated by passaging with cell dissociation buffer allowing one plate to be used for DNA analysis. CRISPR screening was performed by PCR. Homology directed integration of the HDR template introduced a unique Eag1 restriction site enabling a simple and rapid 96 well plate-based PCR screen. Cells were lysed with DNA lysis buffer (QuickExtract, Lucigen) and PCR performed with the primers (Forward TTTTCCCATACCCCTTCGGG, Reverse AGTCATTGGGGAAGGTCTCG), PCR products were digested with Eag1 and analysed by gel electrophoresis. PCR amplicons from candidate edited clones were sequence verified (Eurofins) and sequence analysed using CRISPR-ID software. Absence of editing at Deskgen predicted off-target sites were confirmed by PCR amplicon sequence analysis. Candidate edited clones were identified and expanded in E8-Flex medium.

**hIPC microglial differentiation**

hiPSCs were differentiated to microglia according to (Haenseler et al., 2017). Briefly, hiPSCs treated with 10 µM Y-27632 were dissociated using accutase and resuspended in mTesR medium (Stem Cell Technologies). Embryoid bodies (EB) were formed by aggregating 20,000 cells for 24 hours in 20 µL hanging drops. EBs were collected and grown in suspension in 3TG differentiation medium (mTesR supplemented with 50 ng/mL BMP-4, 20 ng/mL SCF, and 50 ng/mL VEGF-121). These EBs were cultured for 6-8 days (until initiation of EB cyst formation was observed) with half medium change after 2 days. Approximately 20 cystic EBs were plated per well of tissue culture-treated 6-well plates and cultured in 3 ml hematopoietic medium (X-VIVO 15 (Lonza, LZBE02-060F) supplemented with 2 mM GlutaMax, 100 U/ml penicillin, 100 µg/mL streptomycin, 100 ng/ml M-CSF, and 25 ng/ml IL-3) with half medium changes every 3 days. After 2 weeks differentiation myeloid progenitor cells were harvested from the culture supernatant and plated on poly-D-lysine (0.1mg/ml)/fibronectin (0.5 µg/cm2) coated culture plates in microglial differentiation medium (1:1 mix of ADF:ACM media; ADF is Advanced DMEM/F12 DMEM (ThermoFisher), 2% B27 supplement (50X Thermo Fisher Scientific, 17504044), 2 mM GlutaMax, 100 U/ml penicillin, 100 µg/ml streptomycin. ACM is astrocyte conditioned ADF medium; hiPSC-derived astrocytes were differentiated as previously described (Serio et al., 2013). ACM was derived by pooling ADF media from confluent Nunc T500 triple layer flasks (ThermoFisher), ACM aliquots were stored at -80oC. A single ACM batch was used for all experiments.

**Immunocytochemistry assessment of microglia differentiation from hiPSC**

Cultured cells were washed once in PBS and fixed with 4% paraformaldehyde (10 min room temperature) followed by 3 PBS washes. Cells were permeabilised by treating with 100% ice cold EtOH for 2 mins at room temperature followed by 2 x PBS washes. Fixed cells were incubated with blocking buffer (3% (v/v) goat/chicken serum, 0.1% (v/v) Triton-X-100 in PBS) for 1 hour at room temperature before overnight incubation at 4oC with primary antibodies diluted in blocking buffer. Primary antibodies used were anti-IBA-1 (1:100 Abcam AB5076), anti-Glut5 (1:100 R&D systems MAB1349), anti-TMEM119 (1:100 Abcam AB185333) and anti-P2RY12 (1:100 Abcam AB188968), Following overnight incubation, cells were 3 times with PBS prior to a 1 hour incubation at room temperature in the dark with fluorescent secondary antibodies diluted in blocking solution. Secondary antibodies used were Alexa Fluor 594 chicken anti-goat IgG (Invitrogen A21468), Alexa Fluor 594 goat anti-rabbit IgG (Invitrogen A11037) and Alexa Fluor 488 goat anti-mouse IgG (Invitrogen A11029) all at 1:400.

Coverslips were subsequently incubated with Hoechst 33258 (Thermo Fisher Scientific) at 1:5000 in blocking buffer and mounted on microscope slides (immu-mount, Fisher, 9990402). Images were taken using a Leica DM18 confocal microscope.

**RT-qPCR**

RNA was extracted using RNAeasy kit (Qiagen) following manufacturers protocol. All RNA was then quantified and normalised to 500ng. SuperScript IV first strand synthesis kit (Invitrogen) was used to create cDNA and gene expression was analysed using the Quantstudio 5 (Applied Biosystems) and primer/probe assay for murine Plcg2 Mm01242530_m1 and Gapdh Mm99999915_g1 (Life technologies).

**Western blotting**

For MOP cells, cell lysates were prepared using RIPA lysis buffer solution (Santa Cruz sc-24948). Protein extracts (20 µg) were denatured at 70oC for 10 minutes and loaded on a Bolt 4-12 Bis-Tris plus gel (Invitrogen) then transferred to nitrocellulose membrane (Novex). Membranes were blocked for two hours in 5% (w/v) dried milk powder in TBST then exposed overnight to anti-PLCG2 (BioRAD AHP2510) and anti-β-tublin HRP conjugate (Cell Signalling 5346) at $4^{o}$C. Membranes were then washed with TBST and exposed to an anti-rabbit-HRP antibody (Jackson Immunoresearch 711035152) for 2 hours. The membrane was then developed in ECL (Thermofisher 32106) and imaged on a G:Box Syngene using GeneSys software. Bands were analysed using Image ProPremier software.

For hiPSC-derived myeloid precursors whole cell lysates were prepared using ice cold RIPA buffer (Sigma) supplemented with mini protease inhibitor (Roche, 11836170001) and PhosStop reagent (Roche, 049068455001). Protein extracts (20 µg), were denatured in loading buffer at 70oC for 10 minutes and loaded into a 4-12 % NuPAGETM Bis-Tris plus gel (Life Tech, NP0336PK2) and ran in MOPS SDS running buffer at 165V for 45 minutes. Gels were transferred to nitrocellulose membranes that were then blocked with 5% (w/v) skimmed milk and probed with anti-PLCγ2 (Santa Cruz, sc-5283) overnight at 4oC or anti-α-tubulin (loading control, Abcam, ab7291) overnight at 4oC. Membrane were washed in PBS/0.1% Tween and incubated for 1 hour at room temperature with IRDye 800CW Goat (polyclonal) Anti-Mouse IgG (H+L), Highly Cross Adsorbed (1/10,000, LI-COR, 926-32210) prior to washing in PBS/0.1% Tween and imaging on a LI-COR Odessey CLx imaging system. Bands were analysed and quantified using the LI-COR analysis software.

**Oligomerization of Aβ1-42**

Amyloid Beta (Aβ1-42) oligomers were produced in a similar manner to that described in (Ryan et al., 2010). Briefly, Aβ1-42 (A9810 Sigma) was suspended in hexafluoroisopropanol (HFIP, Sigma-Aldrich) to 1 mM. HFIP was then dried with a nitrogen stream and then lyophilized using a Speed-Vac. Samples could then be stored for further use at -20 ºC. Peptide films were resuspended in DMSO to 5 mM. Samples were then sonicated for 10 min, diluted to 200 µM with ice cold PBS + 0.05% SDS, and vortexed for 30 sec. Aggregation was allowed to proceed for 24 hours at 4 °C. The solution was further diluted with PBS to 100 µM and incubated for 2 weeks at 4 °C.

**GapmeR knock down of mouse Plcg2**

Plcg2 knockdown was achieved in M-MØP cells using an antisense LNA GapmeR produced against mouse Plcg2 (Qiagen). Cells were grown to 80% confluence (~2.4 × 105 cells per well) in 6 well plates. 3 µl of Liptoectamine RNAimax (Invitrogen) was diluted in 150 µl RPMI media and mixed with 150 pmol of siRNA in 150 µl RPMI media and left for 5 minutes. 250 µl of siRNA-lipid complex was added to cells in 3 ml RPMI. Cells were left for 48 hours. Cells were then washed and used for Ca2+ imaging (as described below) and knock down of Plcg2 was confirmed with qPCR (as above). ΔΔCt values were used to calculate fold change and comparison of experimental and control samples used to measure knockdown. Negative and transfection control were performed in parallel.

**Live cell DAG Assay in vitro**

Briefly, cells were incubated at 37ºc 5% $CO_2$ for 24 hours with the BacMan sensor, sodium butyrate and receptor control. Cells were then washed with HBSS and left to rest for 30 minutes. Cells were exposed at set time points to anti-FcγRII/III (2.4G2) antibody at 5 µg/ml (Stemcell technologies), LPS (50 ng/ml) (Sigma Aldrich) or oligomers of Aβ1-42 (40 µM). Designated wells were inhibited by pre exposure for 2 hours with Edelfosine (10 µM) (Tocris) or U73122 (5 µM) (Sigma). Other wells were pre exposed to LPS (50 ng/ml) (Sigma Aldrich) or oligomers of Aβ1-42 (40 µM,) for 4 hours). Green fluorescence was measured using the filter on a BMG Flurostar plate reader (BMG labtech) to detect increased fluorescence per well. Similarly cells were imaged using the EVOS FL Auto 2 with the GFP filter at x40 and intensity was measured with Image-Pro Premier software. A BCA assay was run in parallel to normalize. Fluorescence was further normalized by taking the negative control unstained well at 0 mins as 0 A.U.

**Molecular dynamic modelling of PLCγ2 variants**

P522 and R522 proteins were subjected to 300ns of MD. MD was carried out, in triplicate, using the GROMACS package (Pronk et al., 2013) and the Amber03 force field (Case et al., 2010). The structures were boxed and solvated using the GROMCAS module. The molecule was placed in the centre of a cubic box and solvated using TIP3P charge water molecules. Neutralisation of the system was carried out by adding an appropriate number of Cl– ions to the box in the place of water molecules. The particle mesh ewald (PME) method was used to treat long-range electrostatic interactions and a 1.4 nm cut-off was applied to Lennard–Jones interactions. The MD simulations were all carried out in the NPT ensemble, with periodic boundary conditions, at a temperature of 310K, and a pressure of 1 atm. Each simulation was performed using a three-step process: steepest descent energy minimization with a tolerance of 1000 KJ−1 nm−1; a pre-MD run (PR) with 25,000 steps at 0.002 per second per step making a total of 2500 ps; an MD stage run for a total of 300 ns. Root mean square deviation

(RMSD) was monitored along with the total energy, pressure and volume of the simulation to check for stability.

Resulting structures were analysed for flexibility using the g_rmsf and hydrogen bonding using g_hbond (both GROMACS packages (Yang et al., 2015) all proteins were visualised for structural differences using VMD (Humphrey et al., 1996). Further to this prediction of the functional effect and stability analysis was carried out using HoPE (Venselaar et al., 2010). HoPE analyses the impact of a mutation, taking into account structural impact, and contact such as possible hydrogen bonding and ionic interactions. Flexibility, rmsd, energy, pressure and volume distributions were tested for normality using the Anderson–Darling test. All were not normally distributed, and a Mann-Whitney U test was used to determine any statistical differences between wild type and mutated simulations as well as between simulation repeats. This is done using the wilcox test function in the R stats package.
